# Ultrafine mapping of chromosome conformation at hundred basepair resolution reveals regulatory genome architecture

**DOI:** 10.1101/743005

**Authors:** Yizhou Zhu, Yousin Suh

## Abstract

The resolution limit of chromatin conformation capture methodologies (3Cs) has restrained their application in detection of fine-level chromatin structure mediated by cis-regulatory elements (CREs). Here we report two 3C-derived methods, Tri-4C and Tri-HiC, which utilize mult-restriction enzyme digestions for ultrafine mapping of targeted and genome-wide chromatin interaction, respectively, at up to one hundred basepair resolution. Tri-4C identified CRE loop interaction networks and quantifatively revealed their alterations underlying dynamic gene control. Tri-HiC uncovered global fine-gage regulatory interaction networks, identifying > 20-fold more enhancer:promoter (E:P) loops than *in situ* HiC. In addition to vasly improved identification of subkilobase-sized E:P loops, Tri-HiC also uncovered interaction stripes and contact domain insulation from promoters and enhancers, revealing their loop extrusion behaviors resembling the topologically-associated domain (TAD) boundaries. Tri-4C and Tri-HiC provide robust approaches to achieve the high resolution interactome maps required for characterizing fine-gage regulatory chromatin interactions in analysis of development, homeostasis and disease.

## Introduction

Transcriptional regulation the in mammalian genome involves interplay between promoters at the transcription start sites (TSS) and distal cis-regulatory elements (CREs), particularly enhancers. While an ever-increasing number of methodologies have been established to characterize the location and activity of distal CREs, less are available for assessing their physical interactions^1,2^. The chromosome conformation capture (3C) and its derived assays revolutionized the understanding of the chromatin architecture, revealing folding structures known as topologically-associating domains (TADs) bound by looped boundaries harboring orientation-specific CTCF pairs and cohesins. However, recent studies suggest a somewhat limited impact of TAD and boundary loops on gene regulation than initially proposed^3,4^. On the other hand, interactions between CREs have been implicated to regulate gene expression by multiple approaches, including histone quantitative trait loci (QTL) ^5^, enrichment-based loop detection methods such as PLAC-seq and HiChIP^6–8^, and forced chromatin looping ^9^. Revealation of the fine layer structure at sub-TAD level is thus critical to understand gene regulation from the persective of chromatin conformation. However, such microstructures can only be vaguely interrogated by current 3C methods such as 4C and HiC^10,11^ due to their kilobase-range analytical reolutions much below the CRE mapping techniques.

Despite numerous branches of derivatives of 3C being developed, most of them share a conserved principle to map proximity-ligated genome fragments generated by restriction digestion ^2^. Consequently, the detection of contact frequencies is limited at the restriction sites, resulting in a theoretical resolution cap for the methods. Although a 4 bp-cutter restriction enzyme typically used in the current methods yields on average of 256 bp fragments, the static distribuction of restriction sites on the genome results in a significant portion of loci to be consistently underdigested with > 1kb gap intervals (**Fig S1a**). It is likely that small CREs (e.g. 200-300 bp core size for enhancers^12,13^) within these gaps are insufficiently tagged, raising concerns whether their interaction signals can be robustly detected. While alternative digestion approaches, including use of MNase and DNase^14,15^, have been proposed to address such concern, these enzymes can introduce additional bias due to their differential cutting efficiency at nucleosome-free CREs and the rest of genome. Therefore, novel high resolution methods are required for comprehensive interrogation of cis-regulatory interactions.

## Results

### Tri-4C refines genome fragmentation by triple 4-cutter digestion

To overcome the limitations of current methods to comprehensively detect CRE loops, we developed Tri-4C, a novel targeted chromatin conformation capture (4C) method. In Tri-4C, distal chromatin interactions are probed by *in situ* digestion of genomic DNA using three 4 bp-cutter restriction enzymes (REs), DpnII (MboI), Csp6I (CviQI), and NlaIII (**Fig 1a**). *In silico* analysis of the human genome showed that the fragment size of Tri-4C is 1.9-to 5.2-fold shorter than single 4 bp-cutters (**Fig S1a**). The sticky ends are then blunted, allowing free re-ligation of cutting sites generated by three different REs, which dramatically increases ligation complexity. After sonication, the enrichment of contacts at the target viewpoint is achieved by two rounds of nested PCR using two sequential primers in the vicinity of the cutting site. The chromatin contacts are then identified through paired-end sequencing. Similar to UMI-4C^10^, the sonication ends are utilized as unique molecular identifiers (UMI), to generate a PCR bias-free quantitative interaction map. The Tri-4C protocol can be multiplexed, and the procedure can be completed in 3-4 days.

**Figure 1.**
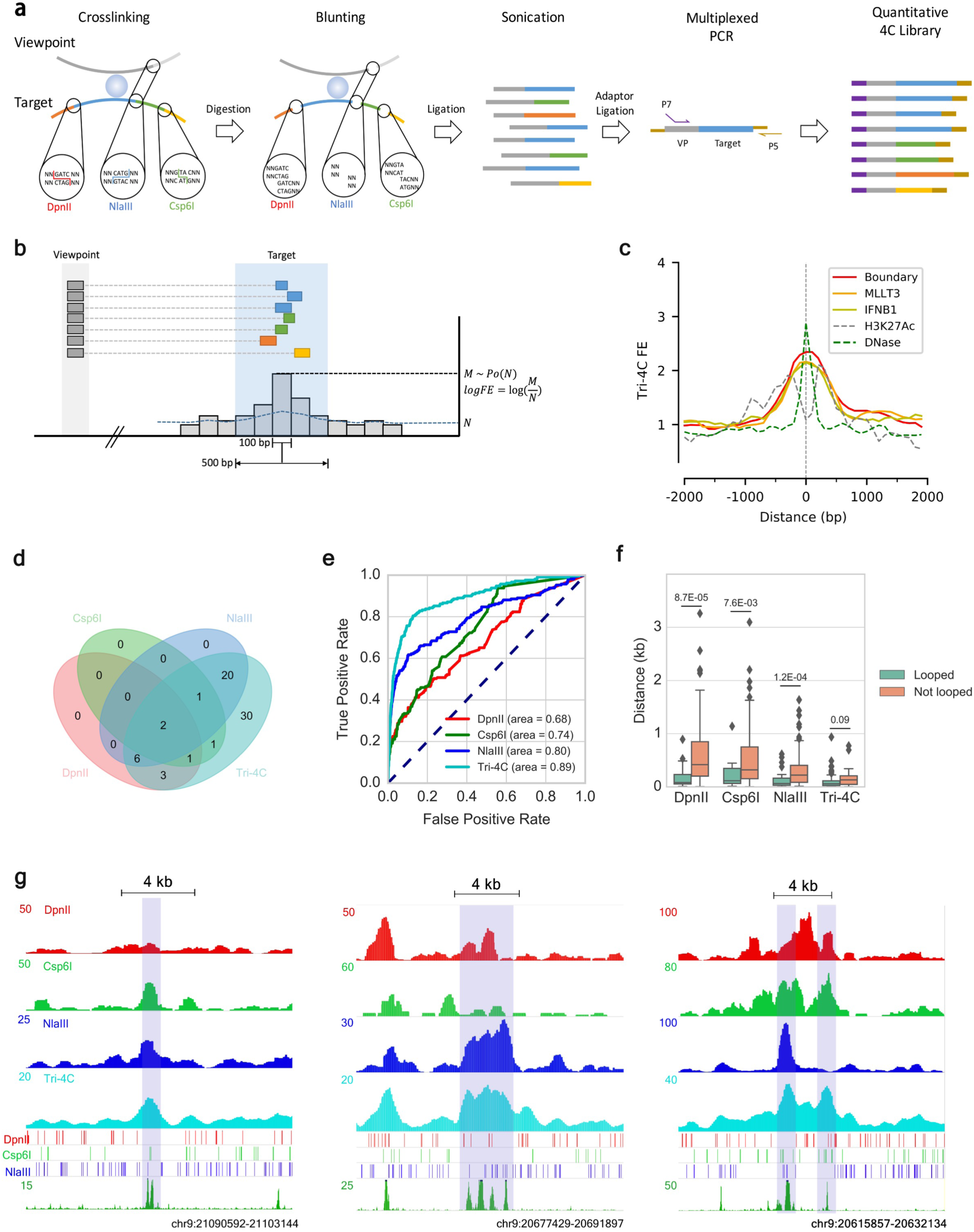
Tri-4C robustly and unbiasedly identifies CRLs. (**a**) Schematics of Tri-4C library construction. (b) Tri-4C loop calling algorithm. Each 100 bp sliding bin collects read count (***M***) from neighboring 500 bp intervals. Significant loops and their loop strengths are determined by Poisson statistics of ***M*** against expected read count ***N*** calculated from the total read counts in the 5-50 kb local backrgound. (**c**) Centerplot of Tri-4C signals at all DHS-marked CREs in the 9p21 IFNB1 TAD. (**d**) Venn diagram of reproducible CRLs (N=2) called for the Boundary viewpoint using Tri-4C and UMI-4C digested by three different restriction enzymes (**e**) ROC analysis using loop signals (−log(p)) for each 100 bp bin steps from Boundary as predictors of intra-TAD DHS peaks. (**f**) Boxplot for distance between intra-TAD DHS peak (N=85) center and the closest restriction site, separated by whether the peak is called to loop with any of the three viewpoints. (**g**) Correlation between raw signals of Tri-4C and UMI-4C and neighboring restriction site patterns at three CRE regions looped with Boundary. Statistic p values were calculated by U test. The y axis of 4C methods denote read count per 10,000 uniquely mapped reads.

To test the performance of Tri-4C, we applied the method to examine the 9p21 interferon B1 (IFNB1) TAD, which harbors 85 putative CREs marked by DNase I hypersensitive sites (DHSs) in IMR90 cells^16^. *In-situ* Hi-C reveals only 8 loops within the TAD that are solely between CTCF binding sites, suggesting the cis-regulatory interactions within the contact domain remain largely elusive (**Fig S2**). We generated the Tri-4C distal interaction profiles on three viewpoints, two promoters (*MLLT3, IFNB1*) and a sub-TAD boundary (Boundary) showing strong CTCF/cohesin binding (**Table S1**, **Fig S2**). To compare with existing techniques, we performed UMI-4Cs (with minor modifications - see Methods) digested individually by DpnII, Csp6I, or NlaIII in parallel on the same viewpoints^10^. To adapt to the multiple ligation ends generated by Tri-4C, we developed a modified pipeline which retained the UMI-recognition property of UMI-4C to remove PCR duplicates for generating quantitative interaction profiles (**Methods**). We found that using the same cell number input, Tri-4C generated on average 5.3-fold more unique contacts than UMI-4C **(Table S2**, **Fig S1b**), suggesting that the detectability of distal interaction proportionally increased with digestion frequency. Consistently, the reproducibility of Tri-4C was significantly higher than UMI-4C, especially at sub-kilobase resolution **(Fig S1c**).

### Comprehensive identification of cis-regulatory loops at hundred basepair resolution

In order to differentiate interaction loops from the local background interactions that occur with high frequency within TADs, we developed an algorithm resembling MACS^17^ to identify loop sites with over-represented interaction read counts. Since 97% of fragments generated by the triple digestion are smaller than 500 bp, we binned reads into 500 bp windows in 100 bp sliding steps, a resolution comparable to the size CREs, and quantified their enrichment against a local background within a 5-50 kb dynamic range (**Fig 1b**, **Fig S1a**). We applied the algorithm to Tri-4C, yielding 233, 138, and 21 reproducible intra-TAD loops, respectively, for the *MLLT3*, Boundary, and *IFNB1* viewpoints. These loops significantly overlapped with a total of 70 CREs marked by DHS, 37 of which were also marked by H3K27Ac (**Fig S3**), which achieved a 4.6-fold increase compared to UMI-4Cs (**Fig 1d**, **Fig S6a**). The loop score of Tri-4C more accurately predicted the positions of DHS-marked CREs and H3K27Ac-marked enhancers than UMI-4C, suggesting that its higher detection sensitivity of cis-regulaotry loops (CRLs) was not compromised by specificity (**Fig 1e**, **Fig S6b,c**). We also examined the mappability, GC content, and restriction site density around the identified loops, finding that loop calling was not significantly affected by these considerations (**Fig S3c-e)**.

The 500 bp resolution (bin size) we chose to perform loop calling for Tri-4C was significantly higher than that with UMI-4C (3-5 kb) or Hi-C/HiChIP (5 kb). To test the impact of higher resolution on CRL detection, we re-analyzed the Tri-4C data with a larger bin size (3000 bp), comparable to previous methods^10,11,18^. At 3 kb resolution, Tri-4C identified on average 35% of the CRLs found at 500 bp resolution (**Fig S4**), with lower signal-to-noise ratios at the overlapping loops, and produced merged loop signals between closely located CREs. Consistently, the 500 bp resolution analysis revealed that CRLs were less than 1 kb long, with the pinnacle precisely aligning with DHS peaks (**Fig 1c**). Hence, sub-kilobase resolution mapping was essential to prevent excess convolution with background, robustly identifying CRLs.

We compared the Tri-4C loop caller with the UMI-4C and the 1D adaptation of *in-situ* Hi-C algorithms, both of which estimate background interactions by using distance modeling based on global interaction profiling (Methods). Receiver operating characteristic (ROC) analysis at 100 bp resolution showed that Tri-4C loops were a strong predictor of DHS-marked CREs regardless of the algorithm used, while loop scores determined by the Tri-4C caller showed the highest accuracy (**Fig S5a**). Furthermore, the CRL strengths (fold-enrichment against background) determined by the Tri-4C algorithm were distance-independent and strongly correlated between viewpoints (r=0.82 between Boundary and *MLLT3*). The correlations obtained by the Hi-C and UMI-4C algorithms were less significant (r=0.48 and 0.29, respectively), probably because of their tendency to over-correct for the distance (**Fig S5b-d**).

### Tri-4C rescues detection of loop signals missed by lack of restriction sites

The UMI-4C profiles generated by three different 4 bp-cutters revealed poorly overlapped subsets of the CRLs identified by Tri-4C (intersection over union < 0.2), suggesting the detection of CRLs by UMI-4C was selective and enzyme-dependent. We case-studied loops that were identified by at least one but not all three UMI-4C profiles, and found that profiels failed to detect the loop unexceptionally lacked restriction sites nearby the CRE (**Fig 1g**). Consistently, the the overall distances between the looped CRE and its nearest restriction site were significantly higher than the unlooped for all UMI-4Cs (**Fig 1f**). Such correlation was not found in Tri-4C, suggesting that its ultrafine digestion of the genome was necessary and sufficient to address the restriction site-depedent loop detection bias. Quantification of loop strength exhibited a negative correlation between the distance between the CRE and the restriction site for UMI-4Cs compared to Tri-4C, suggesting that insufficient genome digestion by single restriction enzyme reduced loop detection sensitivity and dampened their quantification (**Fig S9**).

### Tri-4C loop profile is indicative to cis-regulatory element activities

Investigating Tri-4C-identified loop sites that did not overlap with enhancer marks, including histone modifications and DHS, we found they partially overlapped with ENCODE ChIP-seq signals, suggesting the presence of transcription factor binding sites and therefore regulatory potential (**Fig S7a**). To determine possible regulatory function of these loops, we used CRISPR/Cas9 to delete ~1 kb regions of 4 sites that looped with the *MLLT3* promoter but were devoid of enhancer marks (**Fig 2a**, **Fig S7b, Table S3**). Deletion of two of the sites significantly down-regulated *MLLT3* expression, suggesting that these were *bona fide* enhancers (**Fig 2b**). These functional enhancer loops had escaped detetion by DpnII UMI-4C.

**Figure 2.**
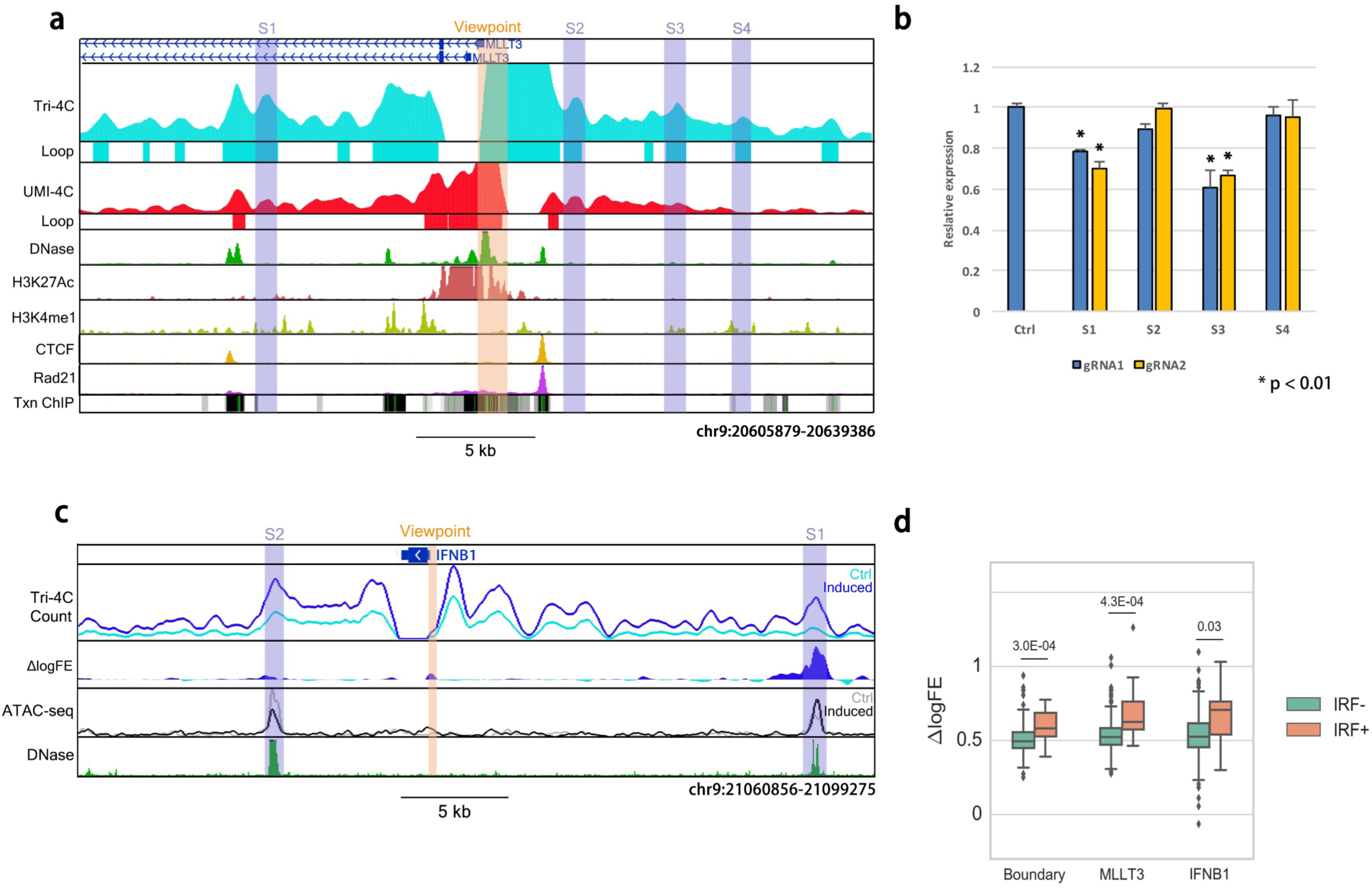
Tri-4C reveals quantitative and functional CRLs. (**a**) Tri-4C, but not DpnII-UMI-4C, indicates looping of MLLT3 with 4 neighboring regions (S1-S4) lacking enhancer marks and CTCF/cohesin. (**b**) Expression of MLLT3 after deletion of these regions using Cas9 and two pairs of guide RNAs (sg1, sg2) quantified by real-time PCR (N=3). (**c**) Alteration of IFNB1 interaction before (Ctrl) and after (Induced) induced expression in IMR90. Top track aligns interaction read count, while second denotes loop strength alteration (ΔlogFE) and shows the loop gain is specific to S1 despite increased count on both S1 and S2. Third track indicates ATAC-seq peak signal of the two enhancers corresponding to the two conditions. S1 is a known enhancer of IFNB1^45^. (**d**) Association between loop strength alterations after IFNB1 induction and IRF(1/2/3/7) motif presence at intra-TAD CREs. Statisitical p values were calculated by U test.

To quantitatively analyze the CRLs called by Tri-4C, we compared the loop strength (i.e. log fold enrichment against local background) with the DHS fold enrichment for all CREs in the locus, finding that they were significantly correlated (**Fig S8a**). Motif analysis indicated that CREs harboring the CTCF motif formed significantly stronger loops with all three viewpoints (**Fig S8b**), consistent with the role of CTCF in mediating chromatin interactions^11,19^. In contrast, this correlation was not revealed when analyzed using UMI-4C.

To test if Tri-4C can reveal the CRL networks underlying dynamic gene control, we induced robust expression of IFNB1 through activation of well-defined antiviral signaling and performed Tri-4C on all three viewpoints^20^. The induction of IFNB1 caused its promoter to interact more frequently with the majority of CREs in the locus (**Fig S10a-c**). However, many of these gains were not significant against the similarly increased local background, and after normalization only 13 CREs showed induced-looping with IFNB1 (**Fig 2c**, **Fig S10d**). The alterations in loop strength with the *IFNB1* promoter significantly correlated with those from the *MLLT3* and Boundary viewpoints, as well as the CRE activities indicated by the ATAC-seq peak strengths (**Fig S10e,f**). The CREs gaining loop strength upon induction were enriched with the motifs of IRF family members, which are key regulators for *IFNB1* activation (**Fig 2d**, **Fig S10f**)^21^. These results indicated that Tri-4C provides the sensitivity and specificity capable of revealing quantitative loop alterations underlying the activities of CREs in the CRL networks.

### Allele-specific Tri-4C quantitively reveals SNP-associated loop alterations

To test whether Tri-4C could differentiate the allelic impact of regulatory variants on CRL networks, we applied Tri-4C to examine the 9p21.3 locus. This locus harbors multiple coronary artery disease (CAD) risk variants, including two functional variants reported to abrogate the function of an enhancer (*ECAD9*) by disrupting TEAD3 and STAT1 binding, thereby misregulating the expression of the target genes, *CDKN2A/B*^22–24^. Using vascular smooth muscle cells (VSMC) derived from a human embryonic stem cell line (H7) that is heterozygous for the risk variants, we performed allele-specific (AS) Tri-4C on ECAD9 (**Fig S11a**)^23^. The AS-Tri-4C profile showed highly cis-specific interaction (>99%), revealing looping of ECAD9 with 25 ATAC-seq-marked CREs in the locus, including both *CDKN2A* and *CDKN2B* promoters (**Fig S11b,c)**. Among the looped CREs, 10 showed differential loop strength between alleles, and in all cases loops on the non-risk alleles were significantly stronger than those of the risk alleles. The stronger loop activity of *ECAD9* on the non-risk allele was consistent with its higher accessibility indicated by ATAC-qPCR (**Fig S11d)**^25^. Lastly, we found that stronger loops were formed between *ECAD9* and CREs harboring TEAD3, STAT1, or SMAD family motifs (**Fig S11e**), consistent with the roles of these factors in regulating *CDKN2A/B*, which are diminished by the CAD risk variants^22,23,26^.

### Tri-HiC maps global chromatin interaction at a hundred basepair resolution with low input

Since Tri-4C has been demonstrated to substantially improve the resolution and CRL detection, we next applied the analogous multi-enzyme approach to develop Tri-HiC for global distal chromatin contact mapping at a hundred basepair resolution. Similar to Tri-4C, the Tri-HiC method utilizes three restriction enzymes to increase genome digestion with the only difference being replacing NlaIII with its isoschizomer, CviAII, to generate 5’ AT overhangs for the biotin labeling (**Fig 3a**). We tested this enzyme combination with Tri-4C on the Boundary and *MLLT3* viewpoints, and found the alternatively digested interaction profiles were highly consistent with Tri-4C using NlaIIl digestion (r = 0.94) (**Fig S12**). To maximize the yield and sequencing efficiency of Tri-HiC, we used Tn5 tagmentation (Illumina), instead of sonication and TA ligation employed in *in situ* HiC, to generate short fragment libraries. Notably, this modification also removed unligated ends due to the insertion nature of transposons. Furthermore, the Tn5 tagmentation step permitted a PCR cycle prior to streptavidin pulldown, which separated the two biotin-labeled strands to increase the capture efficiency for the contacts.

**Figure 3.**
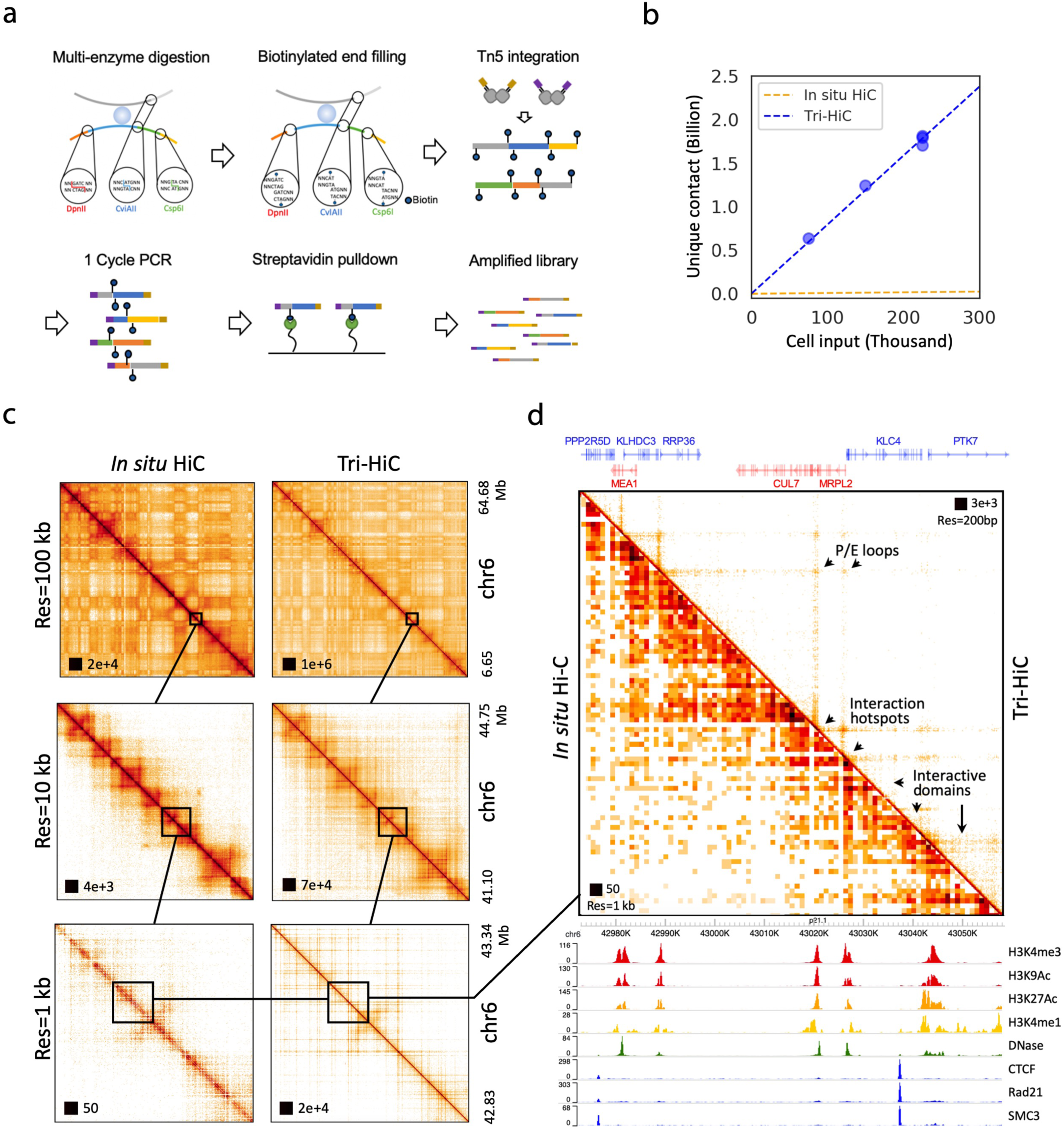
(**a**) Schematics of Tri-HiC library construction. (**b**) Yield curve of Tri-HiC from 5 library replicates in comparison with yield of *in situ* HiC for IMR90 reported by Rao et al., assuming each *in situ* HiC library was constructed with 2 million cells as minimum suggested by authors. (**c**) An example comparison of interaction maps of a locus on chromosome 6 between Tri-HiC and *in situ* HiC at multiple resolutions. (**d**) A 90 kb zoomed-in region from (**c**) with gene and epigenetic annotations. Arrows highlight micro-structural features uniquely revealed by Tri-HiC, including interaction hotspots (anchors for non-specific interaction stripes), promoter-enhancer (P/E) loops, and interaction micro-domains.

We generated Tri-HiC libraries for 5 biological replicates for IMR90 with a total sequencing coverage of 13 billion raw read counts (**Table S4**, Methods). With 80,000-240,000 cell inputs for each replicate, we obtained ~0.6 to 1.8 billion unique contacts, scaling linearly with the input (**Fig 3b**), and 7.2 billion unique contacts in total for all replicates combined. This yield efficiency is about 70 fold more efficient than *in situ* HiC, which typically obtains a few hundred million contacts from 2-5 million cells^11^. Reproducibility tests indicated that Tri-HiC maintained high scores (Pearson R > 0.85) across replicates at a range of resolution up to 500 bp (**Fig S13a**).

Analysis of contact frequency against genomic distance indicated their overall log-linear negative correlation (**Fig S13b**). We observed an up to 10 fold overrepresentation of out-out contacts in the 100 bp −10 kb range, compared to the in-out and out-in pattern, which could indicate that circularized ligation between two ends of the fragment is favored in short distance. Interestingly, the out-out frequency displayed a distinct waving pattern at subkilobase range with peaks at 200, 600, and 1000 bp, which could indicate stable short range structures associated with nucleosome organizations.

We used HiC pipelines Juicer^27^ and Distiller^28^ to map the Tri-HiC libraries with resolution settings up to 100 bp. To evaluate the results, we compared the interaction profiles with the public *in situ* HiC data for IMR90 (1.1 billion contacts)^11^. At lower resolutions (10-100 kb), Tri-HiC recapitulated the macro-structures such as chromatin compartments and TADs revealed by i*n situ* HiC (**Fig 3c**). At 1 kb and sub-kilobase resolutions, however, Tri-HiC uniquely identified multiple microstructures which that were not visible in i*n situ* HiC, including the non-specific interaction stripes extended from distal interaction hotspots, the sub-TAD loops, and interactive micro-domains with sizes as small as a few kb (**Fig 3d**). Contrary to the CTCF/cohesin-centric chromatin architecture revealed by *in situ* HiC, these microstructures were aligned to non-CTCF/cohesin CREs such as promoters and enhancers, revealing their essential roles in organizing the sub-TAD chromatin architecture. Notably, these structures are consistent with the fine-gage interactions recently reported by Micro-C, a high resolution HiC technique utilizing MNase as an alternative approach to overcome the restriction digestion size limit^15^. However, Tri-HiC revealed these structures with substantially lower cell input without digestion bias of MNase at nucleosome-free CRE regions.

To confirm that the identification of the novel CRE-associated structures is not due to our ultrahigh sequencing depth, we specifically compared *in situ* HiC with a Tri-HiC replicate with similar coverage (1.2 billion contacts). In all loci examined, we found that although with some resolution compromise, the low-depth profile recapitulated the structurual features revealed by the pooled library (**Fig S14**). We also noticed that the same 1 kb resolution was sufficient to identify the CRE-associated structures by Tri-HiC but not *in situ* HiC, suggesting that the improvement was not due to the arbitrarily defined resolution bin size but rather can be attributed to the increased detection of interactions, analogous to the advanteges of Tri-4C.

### Genome-wide robust identification of cis-regulatory loops

To identify significant loop interactions, we applied the HiCCUPS^27^ algorithm to Tri-HiC at multiple resolutions. A highest number of 49,619 and 219,399 loops were identified respectively from the 1.2 B contact subset and the 7.2 B pooled library at 2 kb and 1 kb resolution (**Fig 4a**). These numbers were respectively 6.2- and 27.2-fold higher than 8,040 reported by *in situ* HiC^11^. Notably, the loop detection sensitivity of *in situ* HiC sharply decreased from 10 kb to 5 kb resolution, while the power of Tri-HiC peaked at 1-2 kb, suggesting the improvement of loop detection at high resolution was much higher than the overall count ratios. Overlapping the loop profiles between the two assays showed that Tri-HiC reproduced 98% of *in situ* HiC loops at 5 kb resolution (**Fig 4b**). Interestingly, 3,716 in situ HiC loops were only found at 5 kb but not 1 kb resolution, which could indicate limitations of loop detection power at high resolution, possibly due to coverage insufficiency.

**Figure 4.**
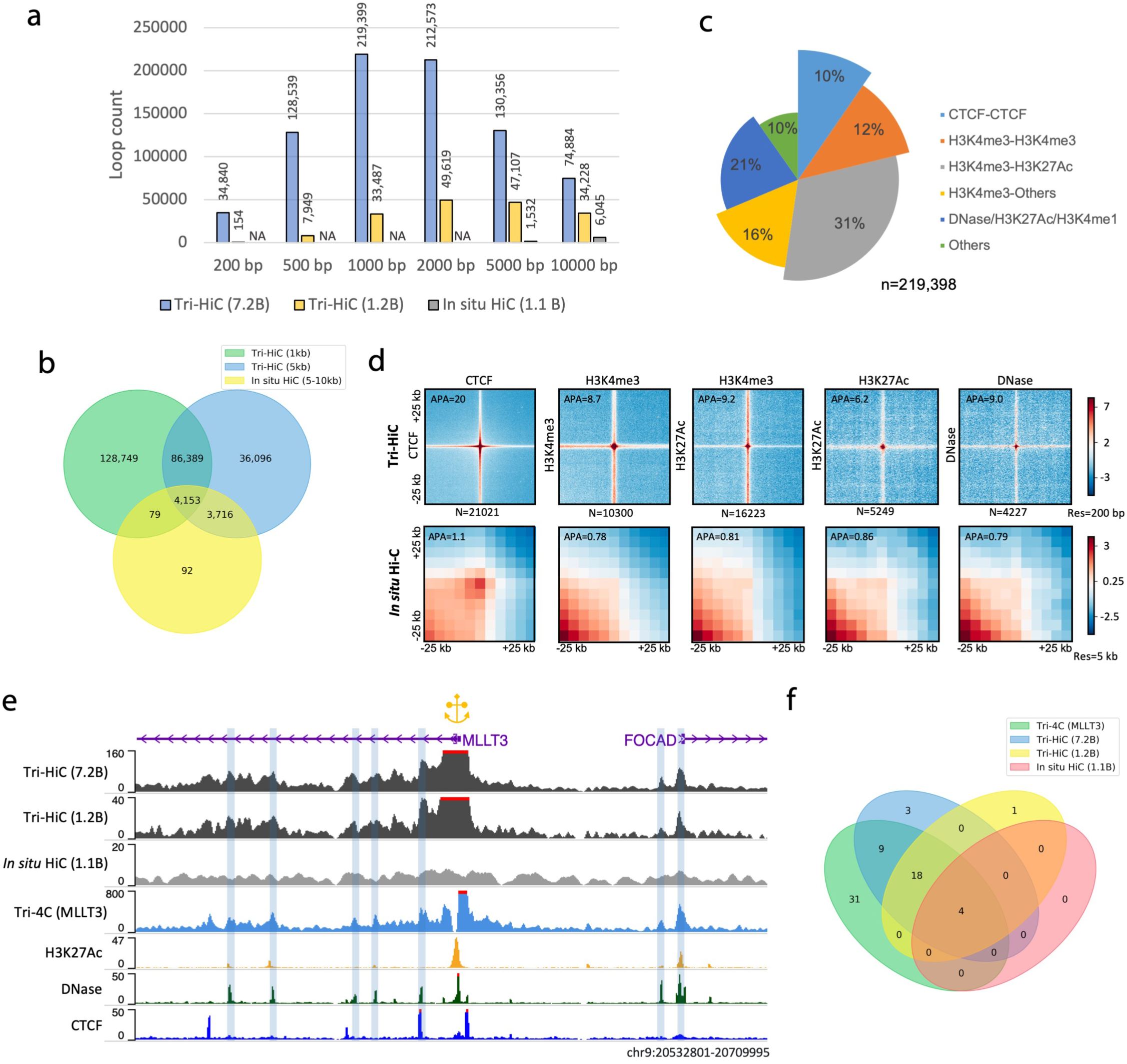
Identification of cis-regulatory loops by Tri-HiC. (**a**) Number of loops called by HICCUPS for Tri-HiC all (7.2 billion contacts), replicated #2 (1.2 billion contacts), and *in situ HiC* (1.1 billion contacts) at different resolutions for IMR90 cells. (**b**) Venn diagram indicating the overlap statistics of loops identified by Tri-HiC at 1 kb and 5 kb resolutions and in situ Hi-C. (**c**) Annotation of loops identified at 1 kb resolution. Loop annotated with multiple categories are classified to the category with the largest bar size. (**d**) Z-score transformed aggregated peak analysis (APA) of loops identified by Tri-HiC at 1 kb resolution sorted by annotations. The aggregated heatmaps are visualized in 200 bp resolution for Tri-HiC and 5 kb for *in situ* HiC. Loops annotated into multiple categories are included only once into the leftmost figure. (**e**) Comparison virtual 4Cs derived from Tri-HiC and *in situ* HiC with Tri-4C for MLLT3. (**f**) Overlaps of MLLT3 intra-TAD DNase loops identified by different methods.

We next annotated the 1kb loops identified by Tri-HiC. The highest fraction of loops (31%) were found between promoters (H3K4me3) and active enhancers (H3K27Ac); by contrast, only 10% of the loops were bound by CTCF sites (**Fig 4c**). This composistion is distinct to *in situ* HiC, for which the majority of loops (58%) were identified between CTCFs (**Fig S15a**). The vast majority (90%) of Tri-HiC loops had at least one anchor overlapped with active CREs, indicating their central roles in distal loop formation. Aggregate peak analysis (APA)^27^ showed that loop strength fold enrichment for Tri-HiC was highest (20-fold) for CTCF loops and lowest for active enhancer loops (6.2 fold) (**Fig 4d**). These numbers were substantially higher than typically obtained from *in situ* HiC (1-4 fold)^11^, with the APA for the same loops of *in situ* HiC showing only positive, but weak, loop signal (1.1 fold) for CTCF loops and no loop signals for all other CRLs. At 200 bp resolution, we observed that the average loop size was typically about 1×1 kb^2^, suggesting that sub-kilobase resolution is required for characterizing the loops without convoluting their signals with the background.

To evaluate the sensitivity of Tri-HiC CRL detection, we compared the virtual 4C profile of MLLT3 from the method with Tri-4C (**Fig 4e**). Out of 62 intra-TAD CRLs detected by Tri-4C, Tri-HiC identified 31 (50%) compared to 4 (6%) by *in situ* HiC (**Fig 4f**). We examined 6 additional promoters located around multiple enhancers, and found that the virtual 4C peaks generally well overlapped with DNase and H3K27Ac peaks, in contrast to the flat signals from *in situ* HiC (**Fig S15b**). Taken together, the improved resolution of Tri-HiC proved to substantially increase the detection of loops mediated by all types of active CREs.

### Revealtion of high non-specific distal interactivity of regulatory elements

Tri-HiC marked loci that formed strong non-specific interactions, visualized as stripes which extended up to hundreds-thousands kb (**Fig 5a**). These stripes resulted in significant overrepresentition of distal interaction read counts, which we could identify using an algorithm similar to the loop caller for Tri-4C (**Methods**). With a stringent Bonferroni threshold (p < 3E-7), we identified 230,720 significant distal interaction hotspots at 100 bp resolution. Annotation of these hotspots indicated their large overlap with all active CRE epigenetic markers (**Fig 5b**), with the highest being 12.2 fold enrichment with DNase, suggesting that high distal interaction activity is a general property of active CREs. Consistent with this interpretation, the distal interactivity score was strongly predictive to all active CRE markers, with the highest AUC score found with H3K4me3 (0.84), CTCF (0.84), and DNase (0.82) (**Fig 5c**). Quantitative comparisons of the interactivity scores among different CREs showed that promoters regions (H3K4me3) were the strongest distal-interacting elements, followed by CTCF binding sites, and both were higher than that observed active enhancers (marked by H3K27Ac) and other DNase-marked CREs (**Fig 4d**). We further compared the motif enrichment between interaction hotspots and DNase peaks, and found that DNase peaks were specifically more enriched with AP-1 class motifs, which have been reported to be depleted in promoters ^29^. In contrast, Tri-HiC interacting regions were more enriched with several CG-rich motifs, including SP1/2, EGR2, and KLF4 (**Fig 5e**). These results together indicated that promoters were a highly-enriched distal interactive element among all CREs.

**Figure 5.**
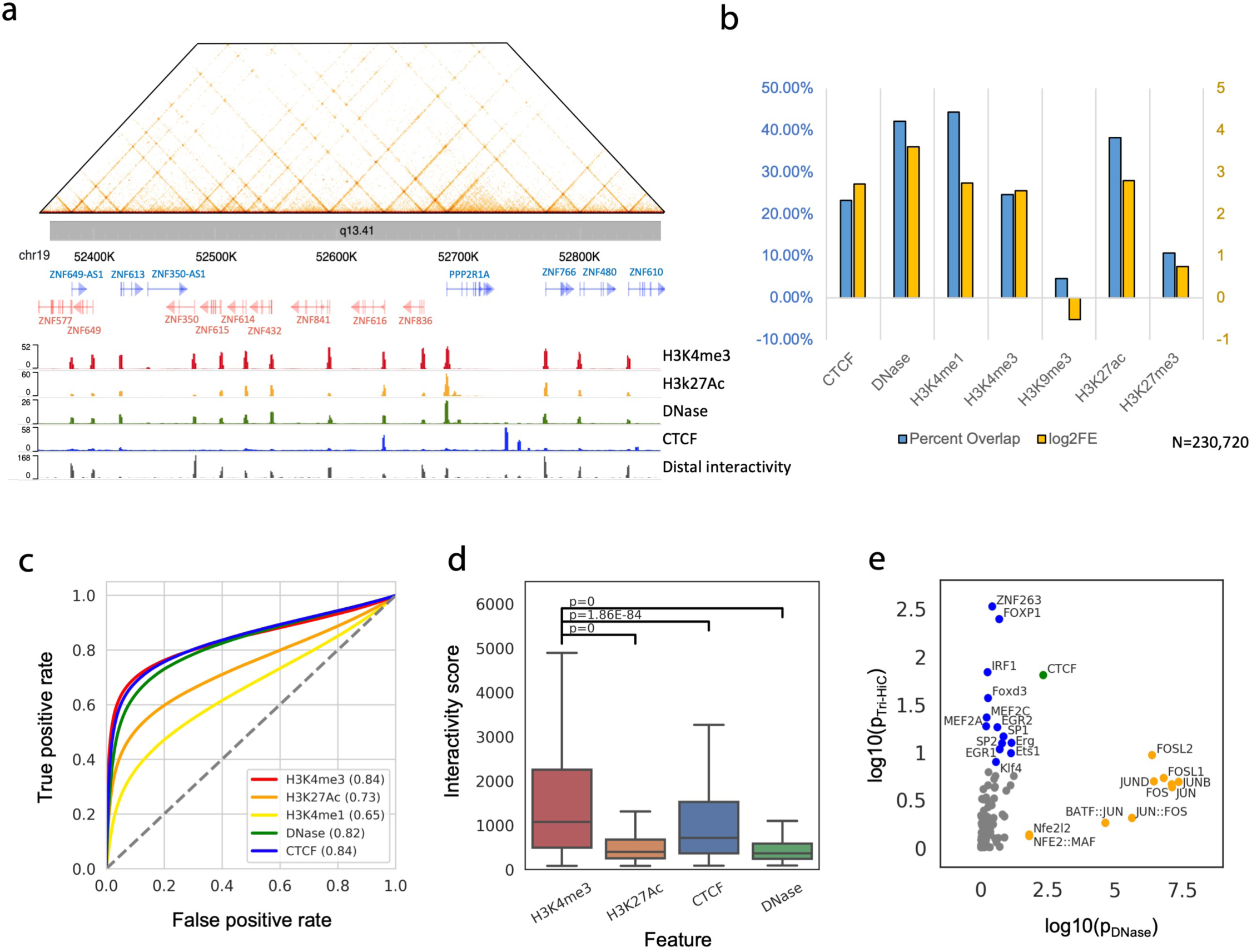
Tri-HiC identifies regulatory elements as distal interaction hotspots. (**a**) An example of gene promoters forming bidirectional non-specific interaction stripes in chr19. Aligned with the landscapes of epigenetic marks, the distal interactivity track at bottom indicates −log10 p value of distal interaction read count enrichment (see Methods). (**b**) Percentage overlap and log2 fold enrichment (log2FE) of distal interaction hotspots identified by Tri-HiC with other 1D epigenetic marks. The interaction hotspots annotated into multiple categories are only included in the leftmost category. (**c**) ROC analysis using distal interactivity scores (−log(p)) for each 100 bp bin steps as predictors of CRE peaks. (**d**) Comparisons of distal interactivity scores for hotspots annotated with different epigenetic marks. Significance of differential interaction scores was calculated by using two-side t test. (**e**) Comparisons of motif enrichment scores (log10 p value) between IMR90 DNase peaks and distal interaction hotspots identified by Tri-HiC.

Interaction hotspots were depleted in heterochromatin marked by H3K9me3, but instead enriched in H3K27me3-marked polycomb-repressed regions (**Fig 5b**). Consistently, we observed interaction stripes from polycomb-repressed genes with strikingly long widths typically greater than 10 kb, suggesting that polycomb repression failed to prevent distal chromatin interactions (**Fig S16**). Notably, despite their extended widths, which allowed visualization at 2-20 kb resolution range in Tri-HiC, these super stripes were not detected by *in situ* HiC.

### Quantitative analysis dissects correlation between loop and stripe formation

Annotation of Tri-HiC loop anchors with distal interactivity indicated that the majority of loop anchors were also interaction hotspots (**Fig S15c**), which is consistent with our findings that CREs often loop with multiple partners within the stripe range, forming interaction networks such as the promoter network in the chr19 KRAB zinc finger protein (KZFP) cluster shown in **Fig 4a**. Such observation raised the question whether the loops identified by Tri-HiC are indicating a stable loop structure between CREs that has been widely referred in the enhancer-mediated transcription mechanisms^30^, or merely a high incidence of co-localization between loci with high distal interactivies. The two scenarios can be distinguished by quantitative analysis to determine whether the interaction frequencies observed at loops are higher than expected from the non-specific interactions from the hotspots. By comparing the loop strengths to the product of the neighboring stripe strengths from the two loop anchors, we found that although they were significantly correlated (R=0.67, **Fig S17a**), the loop strength was consistently higher for all types of CRLs, with the sole exception being promoter-promoter interactions (**Fig S17b**).

To further undertand factors contributing to the CRLs, we deduced the loop strengths contributed from the stripe fold enrichments, and associated the residual loop strength with the motif component on the loop anchors. Hierachial clustering of motifs showed that TFs in several famiies, including STAT, ETS, E-box, and AP-1, interacted stronger with the same family member and tended to have similar interaction preferences (**Fig S17d**). On the other hand, the CpG-associated motifs, such as ZBTB33, NRF1, and E2F, were associated with weaker strength regardless of the loop partner. Importantly, we found that the majority of motifs formed stronger interactions when paired with themselves (**Fig S17c**), which could implicate a general homotypic interaction of TF-bound regulatory regions. Although further experiments are required to clearly interpret these observations, Tri-HiC suggests that homotypic TF pairing is likely to play a significant impact on the loop strength.

### Insulator properties associated with cis-regulatory elements

The Tri-HiC interaction heatmap suggested non-CTCF-bound CREs segregate TADs into further micro-domains (**Fig 3a**). Consistenlty, aggregative analyses showed insulation of background interactions at active CREs similar to CTCF (**Fig 6a**). This finding was further confirmed by insulation score analysis^31^, which indicated precise alignments between boundary signal peaks and interaction hotspots (**Fig 6b**). Quantification of insulation scores showed comparable values between CTCF and promoter regions (H3K4me3), which were both significantly higher than active enhancers (H3K27Ac) and other CREs (DNase) (**Fig 6c**). These results suggested that while all CREs contributed to chromatin domain segregation, promoter regions exerted particularly strong insulation/retention effects that was comparable to CTCF sites typically found at TAD boundaries.

**Figure 6.**
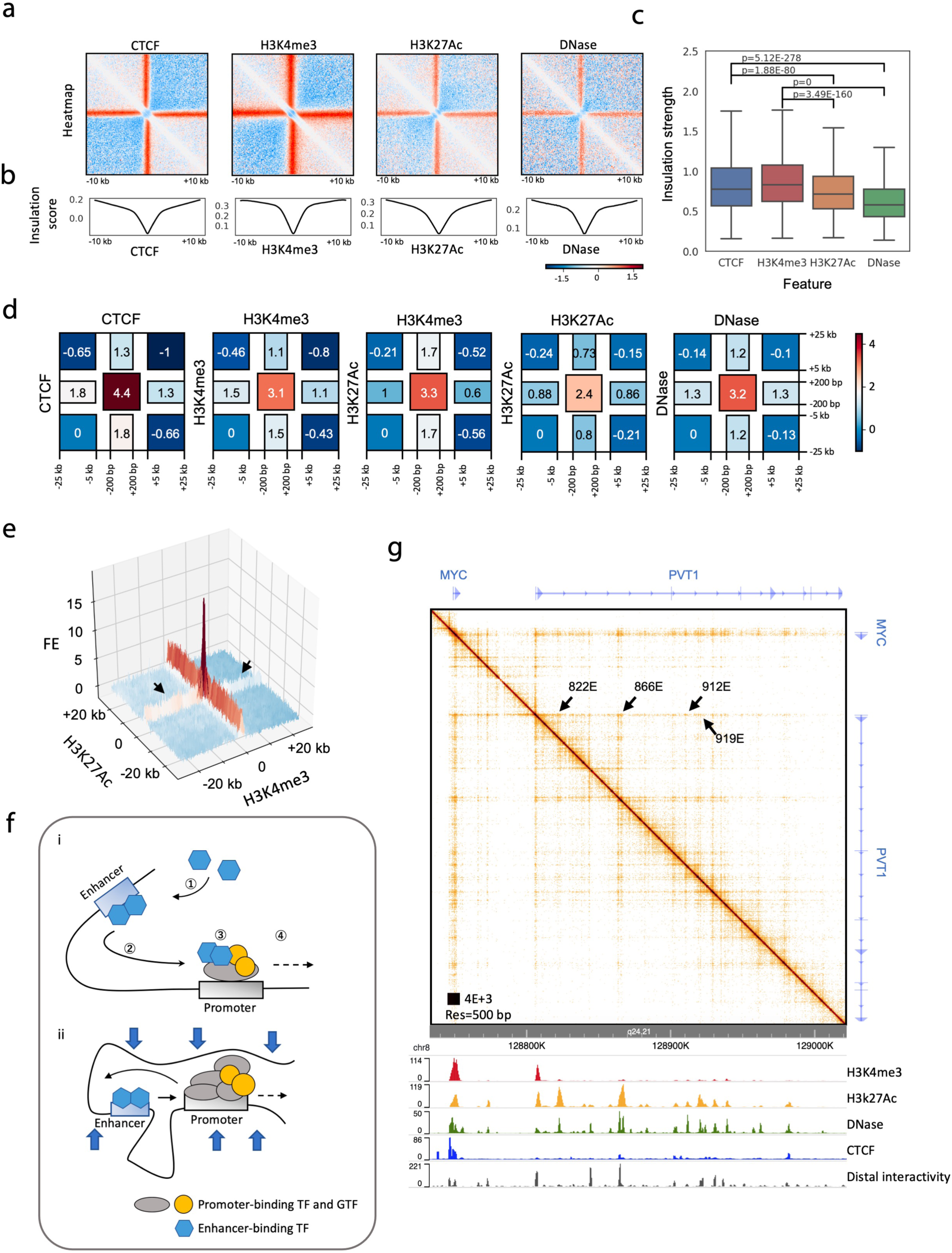
Tri-HiC reveals chromatin insulation associated with cis-regulatory elements. (**a**) Distance-normalized and Z score-transformed piled up heatmaps of distal interaction hotspots annotated with indicated epigenetic marks at 100 bp resolution. (**b**) Piled-up insulation scores for regions indicated in (**a**). Comparison of insulation strength of interaction hotspots, defined as the range of insulation scores within their 20 kb neighboring regions, annotated with indicated marks. Regions annotated into multiple categories are included only once into the leftmost category. (**d**) Relative log2 interaction intensities of the four corners, of interaction loops with indicated annotations. (**e**) A 3D pile-up showing the asymmetry of promoter-active enhancer loops. Black arrows indicate reduction of enhancer (marked by H3K27Ac) stripe strengths after encountering the looped promoter (marked by H3K4me3). (**f**) Hypothetical models interpreting the asymmetric insulation of promoter-enhancer loops. (i) In the “transfer”/asymmetric loop extrusion model, ① enhancers recruit transcription factors and co-factors which ② mediates non-specific interaction stripes before encountering the promoter. ③ Upon the contact, these factors are partially transferred from the enhancer to the promoter, ④ thereby weakening the enhancer interaction stripe downstream. (ii) in the E:P retention model, enhancers are retained after meeting the preferred promoter. The large scaffold complex at the promoter thus blocks the enhancer from walking further downstream, while the TF complex on the enhancer is not sufficiently large to prevent promoter from interacting with the upstream. (**g**) Tri-HiC interaction heatmap of MYC-PVT1 locus in IMR90. Arrows indicate diminishment of interaction stripes of 4 enhancers upon interacting with PVT1 promoter.

To assess the impact of micro-domains on distal interactions between CREs, we analyzed the interaction changes around CRLs by quantifying the intensities of the corners and stripes inside and outside the loops. We found that similar to their effect on background, the promoters and CTCF-bound regions were insulative to their loop partners, reducing their stripe extension to the exterior of the loop (**Fig 6d**). By contrast, CREs such as active enhancers did not prevent the loop partner from interacting further downstream, although a weak insulation of the background was observed. Thus, the differential insulation property of promoters vs. enhancers resulted in their asymmetric loop structure, where promoters could be construed “block” the enhancer from farther distal interactions but not vice versa (**Fig 6e**), consistent with ideas that fine-gage E:P interactions are dynamically favored at the expense of other distal E:P pairing in the same contact domain^32^.

## Discussion

We developed Tri-4C and Tri-HiC to further address the resolution limit of current 3C methods. We demonstrated that the usage of a combination of multiple restriction enzymes was essential and sufficient to capture interactions missed by the currently-existing methods, thereby allowing comprehensive and quantitative identification of cis-regulatory loops at a one hundred basepair resolution. The approach of utilizing high resolution analysis to obtain the true signal-to-background ratio of CRLs is distinct from approaches based on immunoprecipitation enrichment such as HiChIP and PLAC-seq ^8,18^, and avoids inevitable biasis introduced by all selection strategies. Indeed, we expect the principle of multi-digestion would actually be applicable to these methods to further enhance their resolution and loop detectability. Notably, the multi-digestion also improved the contact yield efficiency, reducing the input DNA for Tri-HiC down to hundred nanogram range, allowing affordable Tn5 tagmentation strategy to further lower the amount of input. Despite the known mild GC preference^33,34^, Tn5 has been widely used to uniformly fragmentize genomic DNA for low input high throughput sequencing. Potential tagmentation bias can be later addressed with normalization against the self re-ligation contacts, which accounts for the majority of the Tri-HiC contacts **(Fig S13b**), to avoid false calling of significant distal interactions. These improvements together enabled Tri-HiC to be performed with as few as 0.1 million cells. Such low input requirement should broaden the applicability of the method to samples with limited availabilities, extending its potential clinical utility.

With improved resolution, we found that CREs played central roles in organizing distal chromatin interactions, uncovering finer chromatin structural events underlying the TAD structures implicated by Hi-C assays. Tri-HiC indicated that active CREs formed both non-specific stripes with neighboring regions and loops with other CREs, which corresponded with the loops identified by Tri-4C. These analyses suggested that direct loop interactions with promoters can interpret the transcriptional alterations caused by mutations of distal CREs lacking CTCF and cohesin binding. Our result also suggested that high distal interactivity can be used as a novel parameter to map CREs, providing exemples by using Tri-4C to identify functional regulatory elements. The allele-specific looping detected by Tri-4C, as well as the quantitatve correlation between loop strength and ATAC-seq peak strength, indicated a correlation between distal interactivity and activities of CREs, which is consistent with recent studies showing linkage between enhancer gain and loop formation during cell differentiation^7,35^. However, revelations of super-stripes at polycomb-repressed genes by Tri-HiC suggested that distal interactvity does not fully depend on the activation properties of the CREs.

Tri-HiC showed that CREs are generally insulative and thereby segregating TADs into smaller micro-domains. At TAD level, the domain formation has been well explained by the loop extrusion model in which the convergent CTCF pairs confined cohesins that mediated high background interaction within the TAD ^36^. However, the lack of CTCF and cohesins at promoters and enhancers suggests distinct factors or mechanisms maintaining the micro-domains maintained by these elements. Because Tri-HiC data suggests that both loop extrusion and homotypic E:E and E:P interactions (**Fig 5d**) are aspects of CRLs, it can be inferred that formation of CRLs should involve both the cohesive factor-mediated chromatin junction and interactions between motif-binding TFs. Tri-4C and Tri-HiC provide valuable tools for future studies to dissect the interplays between regulatory loops and TF interactions between regulatory elments.

Contrast to the symmetric insulation on both ends of TAD boundaries, we found that enhancers were insulated in an asymmetric fashion by their looped promoters. To interpret such observation, we proposed two hypothetical models (**Fig 6f**): in a one-sided loop extrusion model, the P-E loop contact results in the enhancers transferring some of the the interaction-mediating/activating machinery to the promoter upon looping, which is stabilized and reduces their own distal interactivity; or a E:P retention model, enhancers are preferentially retained, likely in a cohesin-dependent fashion, to a preferred promoter in the contact domain, thereby retaining their interactions. Both models infer that promoters can decrease the ability of an enhancer to interact with the less-favored potential promoter targets, which is supported by recent findings that the long non-coding RNA *PVT1 is* a insulator for its upstream gene *MYC*^32^. We found the promoter of *PVT1* “inhibited” the extention of the interaction stripes for all its intronic enhancers, including 4 with validated function, to *MYC*, consistent with the concept of retention of enhancer interactions with a preferred cognate promoter (**Fig 6g**). Interestingly, we identified multiple oncogene regions resembing the *MYC* locus where lncRNAs resided between the oncogene and its looped enhancer clusters (**Fig S18**). However, the insulation effect was only found on active and stripe-forming lncRNA promoters such as LINC01798 in the MEIS1 locus. The preferred promoter thus does not always overlay with enhancer proximity.

## Methods

### Cell Culture

IMR90 cells (ATCC CCL-186) were maintained in EMEM (Corning 10-009-CV) with 10% FBS (GEMINI 100-500). To induce *IFNB1* expression, cells were treated with 20 μM 2’3’-cyclic GMP-AMP (Invivogen tlrl-nacga23), 100 ng/mL IFNγ, and 10 ng/mL TNFα for 24 hours^20,37^. Cells were collected at full confluence for all downstream analyses.

Human embryonic stem cells H7 (WiCell WA07) were maintained in feeder-free E8 system (ThermoFisher A1517001). Differentiation of the ES cells to vascular smooth muscle cells was conducted as previously described^38^. Briefly, cells were plated on vitronectin (ThermoFisher A14700) coated surface at 5-10% density. On day 2, the medium was switched to N2B27 (50% DMEM-F12 + 50% Neurobasal medium + 1x N2 supplement (ThermoFisher 17502048) + 1x B27 supplement (ThermoFisher 17504044)) supplied with 10 μM CHIR-99021 and 25 ng/ml BMP4 to induce mesoderm differentiation. From day 5, cells were incubated in N2B27 medium supplied with 10 ng/ml PDGF-BB (Peprotech 100-14B) and 2 ng/ml Activin A (Peprotech 120-14P) to induce VSMC differentiation. Five days after, the Activin A was retrieved from the medium, and the cells were expanded for two population doublings and collected for analysis.

### RNA isolation and qRT-PCR

RNA was isolated from cells using the PureLink RNA mini kit (Thermo 12183020) according to manufacturer’s instructions. DNase treatment was performed using DNA-free DNase Treatment & Removal Kit (Thermo AM1906). To synthesize cDNA, reverse transcription was performed using SuperScript IV Reverse Transcriptase (Thermo 18091050) with oligo-dT primers following manufacturer’s instructions. Quantitative RT-PCR was performed on the Applied Biosystems StepOne Plus platform. For *IFNB1*, a pre-designed Taqman probe (Hs01077958_s1) was used with TaqMan Universal PCR Master Mix (Thermo 4304437). For *MLLT3*, a custom primer pair (Fw: TTTGTGGAGAAAGTCGTCTTCC; Rev: GAGGTGATTCACTGGTGGATG) was used with PowerUp SYBR Green Master Mix (Thermo A25741). Expression was quantified using the delta CT method normalized to *HPRT1* (Thermo 4326321E).

### Cas9-mediated Gene Editing

Guide RNAs (gRNAs) for the Cas9 endonuclease were selected using the CRISPR design tools from Zhang lab (http://crispr.mit.edu). To generate enhancer deletions in IMR90 cells, the IDT ribonucleoprotein (RNP) system was applied following the protocol provided by the manufacturer. Briefly, synthesized crRNAs were annealed with tracrRNA and incubated with Cas9 V3 (IDT 1081058) at equimolar concentrations. To perform NHEJ-mediated deletion, we transfected 1×10^5^ IMR90 cells with 22 pmol of Cas9 RNP using the Neon electroporation system with resuspension buffer R (ThermoFisher) at 1100V, 30ms, 1 pulse. After 72 hours, cells were collected for genomic DNA and RNA extraction. To measure deletion efficiency, target sites were amplified using the validation primers flanking the deletion region as indicated in **Table S3** and examined using electrophoresis.

### ATAC-seq

ATAC-seq was performed as previously described, with minor modifications^21^. Briefly, 50,000 IMR90 cells were collected and resuspended in 50 μl cold lysis buffer (10 mM Tris-HCl pH 8.0, 10 mM NaCl, 0.2% Igepal CA630 (Sigma)). After incubation on ice for 5 min, cells were centrifuged at 800 rcf for 5 min at 4 °C. The cell pellet was resuspended with 50 μl transposition mix containing 25 μl 2X TD buffer (Illumina FC-121-1030), 3.5 μl Tn5 transposase (Illumina FC-121-1030), and 21.5 μl water. The mixture was incubated at 37 °C for 30 min with gentle rotation, and transposed genomic DNA was recovered using DNA Clean & Concentrator-5 (Zymo Research D4013). The library was amplified using NEBNext high fidelity PCR master mix (M0541) containing 1.25 μM customized Nextera universal (Ad1_noMX) and indexed primer with the cycling condition of 72 °C for 5 min, 10 cycles of 98 °C for 30 s, 63 °C for 10 s, 72 °C for 1 min, and final extension at 72 °C for 5 min. The amplified library was purified using SPRI beads (Beckman B23318). A double size selection was performed with 0.5x/1.8x bead volume to remove amplicons > 1000 bp or < 100 bp. Libraries were subjected to 150 bp pair end sequencing on the Illumina HiSeq platform with an expected read depth of 70 million. The FASTQ data was aligned and analyzed using the pipeline from the Kundaje Lab (Github https://github.com/kundajelab/atac_dnase_pipelines).

For ATAC-dPCR for ECAD9, 20 ng of final library was loaded to Quantstudio 3D digital PCR system (version 2, ThermoFisher) and amplified using Taqman array (rs4977575, ThermoFisher C_27869497_10). Allele-specific signal quantification was performed using the online cloud application provided by the manufacturer.

### Tri-4C and single RE UMI-4C library construction

To generate the preamplificiation library, Tri-4C adapted the *in situ* Hi-C and UMI-4C protocols^7,10^. 10^7^ cells were fixed with 1% (v/v) formaldehyde (Thermo Fisher 28906) in PBS for 10 min at room temperature. Fixation was quenched by adding 2.5M glycine, dropwise, to a 0.2M final concentration and incubating for 5 min at RT. Cells were washed with cold PBS twice and pelleted (300 rcf, 4 min, 4 °C) in 2ml Eppendorf LoBind tubes. Pellets could be immediately used for downstream procedures, or snap-frozen with liquid nitrogen and stored at −80 °C.

To prepare crude nuclei, the cell pellet was resuspended in a premixture of 250 μl cold lysis buffer (10 mM Tris-HCl pH 8.0, 10 mM NaCl, 0.2% Igepal CA630 (Sigma)) and 50 μl protease inhibitors (Sigma P8340). After mixing thoroughly, the suspension was incubated on ice for 15 min and centrifuged (1000 rcf, 5 min, 4 °C). The pellet was washed once with 500 μl of cold lysis buffer and carefully resuspended in 50 μl of 0.5% SDS. The suspension was then incubated in a 62 °C heating block for 7 min, followed by mixing with 145 μl water and 25 μl 10% Triton X-100 (Sigma), and incubated at 37 °C for 15 min for quenching. To carry out triple digestion, the suspension was mixed with 50 μl of buffer G (ThermoFisher), 120 U MboI (DpnII) (Thermo Fisher ER0811, 10 U/μl), 120 U Csp6I (CviQI) (Thermo Fisher ER0211, 10 U/μl), 100 U Hin1II (NlaIII) (Thermo Fisher ER1831, 5 U/μl), and x μl of water, where x is determined to bring the total volume to 500 μl, empirically within a range of 100-150 μl, depending on the pellet size.

The genomic triple digestion can be alternatively performed by using a combination of MboI (Thermo Fisher), Csp6I (Thermo Fisher) and CviAII (NEB) to generate consistent 5’ TA overhangs for other 3C-derived protocols requiring biotin-dA filling. In this case, after Triton quenching, the nuclei suspension was mixed with 50 μl of 10x Custmart buffer (NEB) and 100 U CviAII (NEB), and diluted to 500 μl. The mixture was incubated at 25 °C with rotation for 2 hours, and then 37 °C for two hours or overnight after adding 120 U MboI and 120 U Csp6I.

For Single RE UMI-4C^10^ experiments, the suspension was mixed with 50 μl of buffer R, buffer B, and buffer G, respectively, for digestion using 100 U MboI, 100 U Csp6I, or 100 U Hin1II in 500 μl final volume. All digestions were conducted at 37 °C overnight with rotation.

On the second day, the restriction enzymes were inactivated by incubating at 65 °C for 20 min. After cooling to room temperature, end blunting was performed by adding 3 μl of 10 mM dNTP, 8 μl of DNA polymerase I Klenow (NEB M0210, 5U/μl), and 4 μl of T4 DNA polymerase (NEB M0203, 3U/μl), and incubating at 37 °C for 1 hour with rotation. For blunt end ligation, the suspension was then mixed with 460 μl water, 120 μl T4 ligase buffer (NEB B0202), 100 μl 10% Triton X-100, 6 μl 20 mg/ml BSA (NEB B9000S), and 5 μl of 400 U/μl T4 ligase (NEB M0202S/L), and incubated for 4 hours at room temperature with rotation. The processed nuclei were pelleted (1000 rcf, 5 min, 4 °C) and resuspended in 500 μl 1x T4 ligation buffer supplemented with 50 μl 20 mg/ml proteinase K (Thermo AM2546) and 50 μl 10% SDS, and incubated at 55 °C for 30 min. For de-crosslinking, 60 μl 5M sodium chloride was added and the mixture was incubated at 68 °C for 2 hours. Note that this step can also be prolonged to overnight. Phenol-chloroform extraction was performed to recover DNA as follows: The suspension was washed once with an equal volume of Phenol:Chloroform:Isoamyl alcohol (25:24:1), and once with an equal volume of Chloroform:Isoamyl alcohol (24:1). After phase separation, the aqueous phase was transferred to a new 2 ml LoBind Eppendorf tube, mixed with 60 μl 3M sodium acetate, 1 μl GlycoBlue coprecipitant (Thermo AM9515) and 1.5 ml pure ethanol, and incubated at −80 °C for 15 min. The mixture was centrifuged at max speed at 4 °C for 15 min. The supernatant was carefully removed and precipitated DNA was washed twice with 1 ml cold 70% ethanol. The DNA pellet was air dried for 15 min, and resuspended in 130 μl 10 mM Tris-HCl (pH 8.0) for 1 hour at room temperature, or overnight at 4 °C.

The re-arranged genomic DNA was sonicated to 300-400 bp fragments using a Covaris S2 ultrasonicator. The parameter guidelines from the manufacturer were used, with settings of Intensity (4), Duty cycle (10%), cycles per burst (200), and time (80 sec) as starting points. In general, multiple rounds (typically 2) with the above parameters were run to obtain the desired fragment size peak of 300-400 bp, which was confirmed by Bioanalyzer (Applied Biosystems). The fragmented DNA was double size-selected using 0.40x/1.0x of SPRI beads (Beckman) to remove fragments below 100 bp and above 1000 bp, and eluted in a final volume of 70 μl of 10 mM Tris-HCl. To repair the sonicated fragment ends, the eluent was mixed with 10 μl 10x T4 ligation buffer (NEB), 10 μl 100 mM ATP, 5 μl 10 mM dNTP mix, 4 μl T4 DNA polymerase (NEB), 1 μl DNA polymerase Klenow (NEB), and 5 μl T4 PNK (NEB M0201, 5 U/μl), and incubated for 30 min at room temperature. The repaired DNA was purified by using 1.0x SPRI beads, and eluted in a master mix of 94.5 μl 1x NEB buffer 2 and 0.5 μl 100 mM dATP. After removing the beads, 5 μl of Klenow exo- (NEB M0212S/L, 5U/μl) was added, and the mixture was incubated at 37 °C for 30 min for dA tailing. The processed DNA was purified by using 1.0x SPRI beads, and eluted in 20 μl 10 mM Tris-HCl.

We designed a custom Y-shape adaptor to generate Illumina next-generation sequencing libraries:

Foward: GATCTACACTCTTTCCCTACACGACGCTCTTCCGATC*T
Reverse: /5Phos/GATCGGAAGAGCCATACAGC

The oligos were synthesized using the IDT Ultramer service (Integrated DNA Technologies). The forward and reverse single strand oligos were annealed (95 °C for 5 min, down to 25 °C at 0.1 °C/sec temperature gradient) and prepared at 30 μM stock concentrations. Five μl of adaptor was added to the A-tailed libraries, and mixed with 25 μl blunt/TA ligase master mix (NEB M0367). The mixture was incubated at room temperature for 15 min, and purified with 1.0x SPRI beads. After eluting in 50 μl Tris-HCl, the libraries were purified a second time with 1.0x SPRI beads to completely remove the residual adaptors. The final preamplification libraries were eluted in 100 μl Tris-HCl, and examined by Bioanalyzer (Applied Biosystems) to ensure correct size distributions and absence of unligated adaptors. The size distributions of mature libraries after incorporating adaptors were centered around 500 bp.

To generate the final Tri-4C and single RE UMI-4C libraries, we designed a pair of outer and inner primers, based on the restriction enzyme, for each viewpoint to increase amplification specificity (**Table S1**). For amplification with outer primers, eight 100 μl reactions, each containing 400 μg preamplification library, 2 μM universal primer (AATGATACGGCGACCACCGAGATCTACACTCTTTCCCTACACGACGCTC), 0.5 μM viewpoint-specific outer primer for each multiplexed viewpoint, and 1x SuperFi PCR master mix (Thermo 12358010) with 20% GC enhancer, were amplified with the following conditions: 98 °C for 30s, 14 cycles of 98 °C for 10s, 62 °C for 10s, and 72 °C for 60s, and final extention at 72 °C for 5 min. All primers were synthesized using the IDT Ultramer service (Integrated DNA Technologies). The products were pooled and purified with 1.0x SPRI beads, and amplified with the inner primer pair (Illumina P5 + bait-specific P7 index-attached reverse primer) for 14 cycles using the same conditions. After purification with 1.0x SPRI beads, the products were quantified using the Qubit DNA assay kit (Thermo Q32851), examined by Bioanalyzer (Applied Biosystems), and diluted to 10 nM to be sequenced on Illumina platforms.

We aimed for a read depth of 5 million reads for each viewpoint. For a typical library containing 100,000 unique fragments, this results in 50x coverage. The high coverage is desired as Tri-4C generates more reads than single RE UMI-4C with the same DNA input. In practice, the actual yield varies in multiplexed libraries, possibly due to primer efficiency and off-target amplification. A minimum depth of 1 million reads was required for all our experiments. Sequencing was performed on Illumina platforms (MiSeq/HiSeq) in paired read mode with read lengths of 75-150 bp.

### Tri-HiC library construction

The initial crosslinking and cell pellet preparation procedures for Tri-HiC were same to the Tri-4C protocol, with the exception that each pellet included 80-240k cells. The pelleted cells were treated with 250 μl cold lysis buffer with 50 μl protease inhibitors and washed once with 500 μl of cold lysis buffer. The crude nuclei were permeabilized by resuspending in 200 μl of 0.5% Triton X-100 in lysis buffer and incubating at room temperature for 15 min. To carry out triple digestion, the nulei were then pelleted by centrifugation and resuspended in 250 μl of master mix containing 0.1% SDS, 1x Cutsmart buffer (NEB), 10 μl MboI (NEB R0147S), 5 μl CviQI (NEB R0639S), and 5 μl CviAII (NEB R0640S). The digestion was performed at room temperature for 2 hours, and then 37 °C for 2 hours. The restriction enzymes were then inactivated by incubating at 65 °C for 20 min. After cooling to room temperature, end blunting was performed by adding a 50 μl master mix containing 1.5 μl of 10 mM dCTP, 1.5 μl of 10 mM dCTP, 1.5 μl of 10 mM dCTP, 37.5 μl of 0.4 mM biotin-dATP (ThermoFisher), and 4 μl of DNA polymerase I Klenow (NEB), and incubating at 37 °C for 1 hour with rotation. For blunt end ligation, the suspension was then mixed with 200 μl of master mix containing 50 μl 10x T4 ligase buffer (NEB), 2.5 μl 20 mg/ml BSA, and 5 μl of 400 U/μl T4 ligase (NEB). To maximize the ligation efficiency for the low input Tri-HiC protocol, we alternatated the temperature condition by first incubating the mixture for 2 hours at room temperature,then overnight at 4 °C, and lastly 2 additional hours at room temperature. The processed nuclei were pelleted and resuspended in 500 μl 1x T4 ligation buffer supplemented with 50 μl 20 mg/ml proteinase K and 50 μl 10% SDS, and incubated at 55 °C for 30 min. For de-crosslinking, 60 μl 5M sodium chloride was added and the mixture was incubated at 68 °C for 2 hours or orvernight. The DNA content was then purified by phenol-chloroform extraction, as described in the Tri-4C protocol, and resuspended in 15-50 μl 10 mM Tris-HCl, scaling with the input.

To construct Tri-HiC Illumina library, we performed the following reactions defined in units. Each unit started with 300 ng purified proximity-ligated DNA quantified by Qubit (ThermoFisher), which should be obtained from < 100k starting cells. For tagmentation, one unit of input was diluted to 30 μl and mixed with 50 μl of 2x TD buffer and 20 μl of TDE (Illumina 20034197, Lot 20436911), and incubated at 55 °C for 10 min (Note: the amount of TDE depends on the storage condition of the enzyme. We recommend optimizing the exact amount to obtain fragments peaked at 250-400 bp range). The tagmented DNA was purified with DNA Clean & Concentrator-5 (Zymo Research) and eluted in 30 μl elution buffer. The eluent was mixed with 50 μl 2x NEBNext high fidelity PCR master mix, 1.25 μM ATAC-seq primers, and diluted to 100 μl final volume to be amplified by a 2-cycle PCR with the cycling condition of 72 °C for 5 min, 2 cycles of 98 °C for 10 s, 63 °C for 10 s, 72 °C for 1 min, and final extension at 72 °C for 5 min. Meanwhile, 15 μl of 10mg/mL Streptavidin C1 Dynabeads (ThermoFisher) was prepared by washing 3 times with 200 μl of 2x Binding and washing (B&W) buffer (10 mM Tris-HCl pH 7.5, 1 mM EDTA, 2 M NaCl). The beads were resuspended with 100 μl 2x B&W buffer and were diretly added to the PCR product, mixed with pipetting, and incubated for 15 min at room temperature with rotation. Beads were then separated on a magnet and washed 3 times with 500 μl 1x B&W buffer (5 min incubation each), 1 time with 10 mM Tris-HCl, and finally resuspended in 100 μl PCR solution containing 50 μl 2x NEBNext high fidelity PCR master mix and 0.5 μM ATAC-seq primers. A following PCR reaction was performed with the cycling condition of 98 °C for 30 sec, 8 cycles of 98 °C for 10 s, 63 °C for 10 s, 72 °C for 1 min, and final extension at 72 °C for 5 min (Note: we recommend to remove beads from the reaction at the end of the 4^th^ cycle). The amplified library was size-selected and purified by 0.55-1x SPRI beads before QC and sequencing.

For the Tri-HiC assay of IMR-90 described in the manuscript, 5 biological repeats were performed in starting with 80k, 160k, 240k, 240k, 240k input, which corresponded with 1,2,3,3,3 units of reaction. Final libraries were pooled and sequenced in one single flow cell (expected 10 billion reads output) of the Illumina Nova-Seq S4 platform using the 2×100 bp pair-end sequencing.

### Tri-4C Data Analysis

#### Generation of contact probability map

Due to the utilization of the multiple restriction enzymes, most currently existing C pipelines are not applicable to Tri-4C. However, alignment and processing is straightforward, as described below. After demultiplexing, the reverse end (R2, viewpoint end) of the FASTQ file was used to filter reads that were correctly ligated with the viewpoint by matching the sequence head with the inner primer sequence and the padding sequence. We used FASTX Barcode Splitter (http://hannonlab.cshl.edu/fastx_toolkit/index.html) for this step, with an allowance of 1 mismatch. The tool also trimmed the viewpoint sequence during the process. For allele-specific analysis, the reads were splitted by matching the allele with the tag SNP on the padding sequence using an awk command. The undigested/unligated ratio was calculated at this step by measuring the fraction of trimmed reads, starting with the immediate downstream sequence from the viewpoint. After trimming, the residual reverse end was mapped together with the forward end (R1, sonication end) using BWA mem with default paired end alignment settings against the hg19 genome. The aligned reads were deduplicated according to the mapped position of the sonication end (5’ for the reads on the + strand and 3’ for the - strand) by using a simple AWK script. We considered reads with sonication ends separated by 1 bp as duplicates based on the observation that the Illumina platform occasionally bypassed the first nucleotide of the read. Reads with low quality (MAPQ = 0) were removed from analysis. The complexity of the library (number of unique reads) and intrachromosome ratio were meansured at this stage. Reads from 1kb upstream to 2kb downstream of the viewpoint were removed as these regions were consistently highly interactive and subjected to over deduplication due to saturation of unique sonication ends. Interchromosomal interactions were also excluded from downstream analysis since no loop-like interaction hotspots outside the same chromosome were observed. For standard analysis, the processed reads were binned in 500 bp, with a sliding step of 100 bp for both visualization and other downstream analyses, with the exception of reproducibility tests (**Fig 1c**, **Fig S3c, d**), where reads were binned with the indicated bin size and equal step size. For comparison of the analysis at the conventional lower resolution, reads were binned in 3000 bp, sliding at 100 bp. Of note, we used the entire aligned sequence for this step, instead of assigning each read to its corresponding restriction site, for two reasons. First, we observed a clear directional bias on the target restriction sites, which indicated that the real contact point was not at the RE site but in its close vicinity. Thus, a short overhang during alignment created a benign bias pointing toward the real interaction spot. Second, the Tri-4C method was designed for promoter/enhancer viewpoints which often contained high/low GC contents. The difficulty of designing primers for these regions sometimes results in a long padding sequence and an unmappable short residual sequence on the reverse end after trimming, yielding reads that cannot be matched to the corresponding RE sites. The piled raw read count bedGraphs were used to perform peak calling. To ensure fair comparison, single RE UMI-4C libraries were analyzed in parallel using the same pipeline.

#### Loop Peak Calling

We used a local fold enrichment-based strategy to identify significant interaction loop peaks for the Tri-4C and singe RE UMI-4C data. Thus, the expected number of reads (background) for a given bin with read count *M* was estimated by taking the average of neighboring bins. We used the smallest mean value of 5 kb, 10 kb, 20 kb, and 50 kb intervals centered at the bin location to represent the background *N*. Then, significance *p* values were calculated by *p* = *Pr{X* ≥ *M}* given *X* ~ *Poisson(N)*. Of note, this step can be achieved by feeding the MACS2 bdgcmp function with the background and signal tracks using -m ppois mode^18^.

To identify significant and reproducible peak regions, bins were scored with the −log10(−log10(p)) value, and those with a score > 0 (p < 0.1) in all replicates were collected and analyzed by IDR package^19^ (Github https://github.com/nboley/idr) using the following settings

~~~
--initial-mu 1.5 --initial sigma 0.3 --initial-rho 0.8
~~~

Bins with IDR < 0.05 (score ≥ 540) were considered significant and merged. We defined a minimum length of 300 bp for calling significant distal loop peaks.

The UMI-4C^10^ and 1D adaptation of *in situ* Hi-C^11^ loop calling algorithms were used for comparison. For both algorithms, distance-dependent decay of interaction frequency was calculated at genome-wide level using IMR90 *in situ* Hi-C data at 5 kb resolution. The decay function was further smoothed to 500 bp bins (*W*) in 100 bp step size resolution by using linear interpolation to obtain the *F(d)* (UMI-4C) or *E*\* (Hi-C) suitable for high resolution analysis in Tri-4C. For the UMI-4C algorithm, the background of each Tri-4C profile was obtained by re-allocating the total intra-TAD Tri-4C read counts (*N*) to each bin according to *F(d)*. The enrichment p value of a bin with expected read count of *E* and actual count of *E1* was calculated by fitting to binomial distribution: *Pr*(*B*(*N*, *E*/*N*) > *E*_1_). For the 1D *in situ* Hi-C algorithm, the adjusted expected read count for each bin *E*^*d*\*^_*i*_ can be calculated by the filter fomula

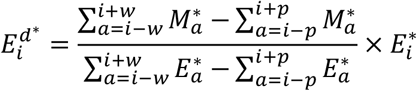

Where p and w were set at 2 and 100 to be corresponding with 500 bp peak and 20 kb background size. This expected count was compared with actual read count *M*_*i*_ using the Poisson statistics. The p values obtained from UMI-4C and Hi-C algorithms were directly compared with Tri-4C raw p value by the CRE ROC analysis.

To investigate for potential artifacts during Tri-4C loop calling due to mapping bias, the GC content and restriction site density under triple enzyme digestion for the locus analyzed by Tri-4C were obtained by directly analyzing the hg19 genome sequence of the region. Mappability of the region was obtained from the ENCODE mappability track available on the USCS genome browser. The average GC content, restriction site density, and mappability for the 10 kb neighboring regions for all loop sites called by Tri-4C were calculated at 100 bp resolution. To generate the background for comparison, a set of 10,000 equal size genomic intervals were randomly selected in the locus. The mean for each set was calculated after removing intervals whose center fell within Tri-4C loops or repeat regions.

#### Data Reproducibility

Each Tri-4C and SRE-4C was performed in two technical replicates. Reproducibility of intrachromosomal interaction was measured by Pearson’s correlation r using the Python numpy.corr function after binning the contacts in the size described in **Fig1C**.

#### Analysis of Interaction Frequency and Loop Strength

For frequency-based analysis, the read count for each bin was converted to interaction frequency by normalizing against (1) total intrachromosomal interactions, which we referred to as normalized interaction or (2) total intrachromosomal interactions of a reference viewpoint in a multiplexed run, which we referred to as relative interaction frequency. The purpose of the latter method was to control for the significant change in total read count generated between different experiental conditions, as in the IFNB1 viewpoint at the baseline control compared to the IFNB1 induced condition (**Fig 2C**, **Supp Fig 9B**). Thus, the reference point was selected based on the standard of exhibiting the least variation in unit read count yield among different conditions, and in our experiment the Boundary viewpoint was chosen for that reason. Subtraction analysis was performed by directly calculating the interaction frequency difference between two tracks in comparison after normalization. A multiplication factor of 10,000 was applied to the normalized interaction frequency to simplify the display when presenting the interaction map on UCSC/WashU track.

For loop-based analysis, loop strength, i.e. log fold enrichment (logFE) was calculated by *logFE* = *log10(M/N)*, where M and N are the actual and expected read count for each sliding window as indicated in the Peak Calling section. This step can be perfomed with MACS2 using -m logFE. A 1.0 pseudo count was given to calculate logFE to resolve zero division as well as attenuating noise level at regions with sparse mapped counts. Differential loop strength was calculated by measuring the logFE difference between two tracks. For presentation in **Fig 2C**, the logFE was weighed by frequency at basal condition.

#### Allele-specific Loop Calling

We used a likelihood ratio test, based on the loop strength, to determine whether a interation loci displayed allelic bias. Specifically, for each 100 bp sliding window we obtained four vectors: *M*_*ref*_, *M*_*alt*_, *N*_*ref*_, and *N*_*alt*_, where *ref* and *alt* denoted the allele genotype, and *M* and *N* denoted the actual and expected read count, as indicated above. Each element in the vector represented one replicate. Firstly, the elements in *N*_*ref*_ and *N*_*alt*_ were normalized to their respective mean, and the scalars were used to normalize their corresponding *M*. Then, we had 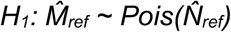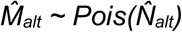, and 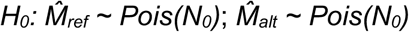, where *N*_*0*_ denoted the mean of 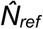 and 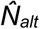. The likelihood ratio *L* was converted to p value by applying the Wilks’ theorem, namely −*2ln(L)* ~ *χ*^*2*^*(1)*, subjected to subsequent Bonferroni correction where n equaled to total number of assayed intervals in the locus.

#### Analysis of Cis-regulatory Element Interaction Network

To annotate CREs and active enhancers, respectively, DNase and H3K27Ac peak position and intensity for IMR90 cells were obtained from the Roadmap Project web portal (https://egg2.wustl.edu/roadmap/web_portal/). For the comparison of loop strength and interaction frequency between viewpoints with DNase peak intensity, linear regression models were built in Python using the scipy.stats.linregress function.

#### Receiver Operating Characteristic (ROC) Analysis

CRE positions, defined by Roadmap DNase peaks, were converted to 100 bp 1/0 tracks, with 1 indicating the presence of peaks inside the bin. The ROC curves were built by using loop scores (−log(p)) obtained from Tri-4C and UMI-4C as predictors for the peak positions in the converted DNase track, using Python sklearn.metrics: roc_curve and auc function with default settings.

#### Motif Analysis

The DNA sequences for all Tri-4C peak regions in both baseline control and IFNB1-induced conditions were extracted. Motif prediction was performed using TFBSTools (R platform) with accession to the JASPAR2018 database^39,40^. A minimum score of 90% was set to discover matched motifs. The Lasso CV model (CV=10, iter=10,000) was applied to all motifs to identify factors correlated with ΔlogFE of Tri-4C peaks during induction. Significance of correlation was determined by F statistic and subject to Bonferroni correction.

#### Hi-C and Topologically-Associated Domain (TAD) Definition

The IFNB1 TAD (chr9:19480000-2120000) was defined by the *in situ* Hi-C data of IMR-90^7^. Hi-C Browser (http://promoter.bx.psu.edu/hi-c/index.html) was used to visualize the Hi-C interactions in the TAD.

#### Code Availability

The Shell pipeline used to align Tri-4C data and interaction loop analysis is available on Github (https://github.com/kimagure/Tri-4C).

### Tri-HiC Data Analysis

#### Contact Map Alignment

The raw Fastq for Tri-HiC was aligned to hg19 genome and processed to obtain the interaction contact matrix in .mcool format by applying the Distiller pipeline ^28^ with default configurations at resolutions of 100, 200, 500, 1000, 2000, 5000, 10000, 20000, 50000, 100000, 250000, 500000, and 1000000 bp. The pairsam intermediate output from Distiller was processed by Juicebox Pre ^27^ to generate contact maps in .hic format with same resolution setup. We kept the alignment with filtering of MAPQ > 0 and > 30. For all analyses downstream we chose MAPQ > 30 by defult, although in all regions reported from this study we did not observe distinguishable difference between the two parameters. For this study, we used HiGlass^41^ to visualize the contact matrices in .mcool format.

#### Reproducibility Test

Reproducibility of Tri-HiC at 500 bp to 20 kb resolution range (**Fig S13a**) was evaluated by the Pearson correlation coefficient between replicates calculated by the HicCorrelate function in the HiCExploer^42^ package. Due to the sparsity of distal contacts at extreme long range which results in mostly empty bins for the high resolutions in our analysis, we restrained the test to 8 Mb intrachromosomal range, which is in line with the distance limit for loop calling (see sections below).

#### Decay Function Analysis

Valid intra-chromosomal contact pairs were binned in 100 bp intervals, and the contact frequency for each bin was calculated by dividing to the total read count. We sorted contact pairs by orientations noted as “in-in (+/−),” “in-out (+/+),” “out-in (−/−),” and “out-out (−/+)”. Smoothed decay curves for each orientation was generated by using the CubicSpline function in scipy.interpolate package from Python.

#### Interaction Hotspot Analysis

According to the decay function analysis (**Fig S13b**), the majority of contacts from Tri-HiC were short range (< 1 kb) in-in self-ligation products, which can be considered as a roughly even background over the genome. Taking the advantage of this property, we defined the interaction hotspots as regions having significant higher fraction of long range contacts, with the short range and long range being respectively defined as smaller and greater than 1.5 kb. For each 100 bp bin, the expected long range read count was calculated by multiplying the short range count with the average long range-to-short range ratio over the 20 kb neighboring background. The statistical significance for the deviation between expected and actual count number was determined by Poisson statistics using the MACS2 bdgcmp -m ppois function^18^. Significant bins with p values smaller than the Bonferroni threshold (3E-7) were merged and considered interaction hotspots. The −log10(p) score from the analysis corresponded to the distal interactivity track displayed in **Fig 4a**, and the average bin score for each peak was reported as its interactivity score for **Fig 4c**.

#### Loop Calling and Strength Analysis

Chromatin interaction loops were called by using the HiCCUPS algorithm from Juicebox^27^. We used the CPU version of HiCCUPS which as described by the developer, searched intrachromosomal loops within 8Mb of the diagonal. Loops were called with the following parameters: -k KR -r 200,500,1000,2000,5000,10000 -f 0.1 -p 5,4,2,2,2,2 -i 10,8,4,4,4,4 -t 0.02,1.5,1.75,2 -d 2000,4000,4000,8000,20000,20000. The same parameters were applied to the public in situ HiC results for IMR-90 at 5 kb and 10 kb resolutions.

The HiCCUPS reported both observed and expected read count from the donut background (expectedDonut) for each loop, and their division defined the signal-to-background fold change of the loop, which we referred to as the loop strengths. Similarly, horizontal and vertical stripe strengths were calculated by dividing the expectedH and expectedV columns, respectively, to the expectedDonut column. The residual loop strength used for motif analysis was defined by the subtraction of log2 horizontal and vertical stripe strengths from log2 loop strength.

The instensities of significant loops identified by HiCCUPS were assessed by aggregate peak analysis (APA) also available from Juicebox with the following parameters: -n 500 -w 125 -r 200 -q 80 -k KR. APA scores from the analysis were directly reported in **Fig 5c**. The intensity matrices obtained from the analysis were transferred to Z-scores and presented in **Fig 5b**. The raw APA matrices were also used to evaluate the relative intensities of corners, stripes, and loops for the insulation analysis in **Fig 6d** by calculating the log 2 mean intensity of each interval indicated in the figure and normalizing against the left-bottom corner.

For the pile-up study at the vicinity of interaction hotspots, the APA was applied with the following parameters: -n 0 -w 250 -r 100 -q 320 -k KR. A distance-based normalization was applied to the obtained matrix by dividing each diagonal to the corresponding diagonal mean from a larger APA obtained with the parameters -n 0 -w 2500 -r 100 -k KR. The resulted matrix was Z-score transformed and shown in **Fig 6a**.

#### Motif Analysis

To annotate the interaction hotspots and loop anchors identified by Tri-HiC, we used the AME^43^ package in the MEME suite to predict motifs in the intervals, with the accession to the JASPAR2018 and TFBSshape databases and default threshold parameters.

#### Insulation Analysis

Genome-wide insulation scores were calculated at 200 bp resolution by using the diamond-insulation function in the cooltools^44^ package with the default parameters. In **Fig 6b**, we piled up the scores within 20 kb region of all interaction hotspots categorized by their annotations. Insulation strengths for each hotspot in **Fig 6c** represented the range (maximum-minimum) of the insulation score of the interval.

### Public Data Usage

The following public datasets for IMR-90 were used for annotations and comparative analysis of Tri-4C and Tri-HiC: DNase (GSE18927), H3K4me1 (GSE16256), H3K4me3 (GSE16256), H3K9me3 (GSE16256), H3K27Ac (GSE16256), H3K27me3 (GSE16256), CTCF (GSE31477), Rad21 (GSE31477), SMC3 (GSE91403), *in situ* HiC (GSE63525), ENCODE Transcription Factor ChIP-seq Clusters (wgEncodeRegTfbsClusteredV3 track).

### Accession codes

Raw and processed data available at NCBI Gene Expression Omnibus, accession numbers GSE119189 and GSE161014.

## Acknowledgments

We thank X. Dong and D. Zheng for discussion on analysis, T. Wang for discussion on statisitcs, and M.G. Rosenfeld and J. Vijgfor their critical reading of the manuscript. Research in the Suh lab was supported by the National Institute of Health (AG017242, GM104459, AG056278, AG057341, AG057433, AG057706, CA180126, HG008153) and by the Glenn Center for the Biology of Human Aging (Paul Glenn Foundation for Medical Research).

## Author Contributions

Y.Z. developed the method, performed experiments and analyzed the data. Y.Z. and Y.S. interpreted the results and wrote the manuscript. Y.S. supervised research.

---

**Figure S1** Tri-4C improves both yield and reproducibility by finer digestion of the genome. (**a**) Distribution of DNA fragment size of human genome digested by indicated restriction enzymes. Numbers on top indicate median and in the parentheses indicate percentage of fragments smaller than 500 bp window size. (**b**) Yield of unique intrachromasomal reads for UMI-4C and Tri-4C on the three viewpoints. (**c**) Reproducibility of interaction profiles binned in 50 kb-50 bp.

**Figure S2** Overview of Tri-4C experimental design at the chromosome 9p21 IFNB1 TAD. Tri-4C profiles of three viewpoints (Boundary, MLLT3, and IFNB1) are displayed under IMR90 *in situ* Hi-C matrix (5kb resolution Rao 2014) obtained from 3D genome browser (Hi-C loops are highlighted with squares). Y axis of Tri-4C tracks denotes interaction frequency multiplied by 10,000. The interaction profiles are aligned with regulatory marks (DNase, H3K4me1) and boundary markers (CTCF, RAD21) for IMR90 cells obtained by the Roadmap Project. Bottom panel shows significant loop interactions between the viewpoints and CREs.

**Figure S3** Tri-4C loop annotation and quality control. (**a**) Overlap between intra-TAD Tri-4C loops and intra-TAD DHS and H3K27ac peaks. (**b**) Overlap of DHS-marked CRLs among the three viewpoints. (**c**) GC content, (**d**) mappability, and (e) restriction site density around regions looped with any of the three viewpoints. Gray background indicates confidence intervals estimated by using 1,000 randomly selected intra-TAD regions not looped with any viewpoints with mappability > 0.5.

**Figure S4** Comparison of Tri-4C profiles analyzed in 500 bp (High res) and 3000 bp (Low res) resolution. (**a**) Overlap of loops falling in CREs. (**b**) Interaction of MLLT3 with neighboring CREs shown by Tri-4C in two resolutions. DHS peaks showing looping at 500 bp resolution are highlighted.

**Figure S5** Comparison of loop calling algorithms. (**a**) ROC analysis using loop scores for each 100 bp bin steps calculated by Tri-4C (Dynamic), 1D Hi-C, and UMi-4C algorithms as predictors of intra-TAD DHS peaks. (**b**) 2D plots comparing loop strength (logFE) on all intra-TAD CREs determined by Tri-4C normalization, or read count using (**b**) UMI-4C or (**c**) Hi-C normalization between viewpoints. Color indicates log distance ratio between the x and y viewpoint (blue = closer to x and red = closer to y). Pearson correlation coefficient r and p value from linear regression are indicated.

**Figure S6** (**a**) Venn diagram of reproducible CRLs (N=2) called for MLLT3 and IFNB1 using Tri-4C and UMI-4C digested by three different restriction enzymes (**b**) ROC analysis using loop scores for each 100 bp bin as predictors of intra-TAD DHS peaks. (**c**) ROC analysis using loop scores for each 100 bp bin as predictors of intra-TAD H3K27Ac peaks.

**Figure S7** Analysis of Tri-4C loops not overlapped with CREs (**a**) Overlap between off-CRE loops with intra-TAD regions showing binding with 5 or more ChIP-seq peaks in the ENCODE combined transcription factor binding track (Txn Factor ChIP V3, 161 factors, all cell lines combined). (**b**) Validation of Cas9 deletion of regions indicated in **Figure 2a**.

**Figure S8** Analysis of Tri-4C loop strength. (**a**) Comparison between loop strength (logFE) from three viewpoints and DHS peak log fold enrichment on all intra-TAD CREs. Pearson correlation coefficient r and p value from linear regression model are indicated. (**b**) Association between loop strength and CTCF motif presence for Tri-4C and UMI-4C based on three enzyme digestion.

**Figure S9** 2D plots between loop strength difference between UMI-4C and Tri-4C (Y axis, ΔlogFE) and the distance between the nearest restriction site and the DHS peak center (X axis, log scale) of all intra-TAD CREs. The p values were calculated by fitting with linear regression.

**Figure S10** CRL alterations after IFNB1 induction. (**a**) Expression profiling of IFNB1 and MLLT3 expression before and after IFNB1 induction by using real-time PCR (N=3, t test). (**b**) Comparison of Tri-4C yield for three viewpoints (N=2) (**c**) 2D plot showing read count of IFNB1 Tri-4C (normalized against Boundary) at all intra-TAD CREs before (X axis) and after (Y axis) induction. (**d**) Venn diagram showing the overlap of CRLs called from IFNB1 before and after induction. (**e**) Comparisons of loop strength alterations (ΔlogFE) between three viewpoints and (**d**) with ATAC-seq peak log fold enrichment changes on all intra-TAD CREs. Pearson correlation coefficient r and p value from linear regression model are indicated.

**Figure S11** Allele-specific Tri-4C for ECAD9. (**a**) Schematics for the allele-specific study design. The viewpoint primer is designed to include a heterozygote flag variant in the padding sequence. Reads are sorted and mapped separately according to the variant genotype. Allele-specific interaction loops are identified by differential loop analysis. (**b**) Allele-specific (AS) Tri-4C profile for ECAD9 in H7-derived VSMCs. Intervals below indicate loop regions called by each allele. Loops on ATAC-seq-marked CREs showing significant allelic bias (FDR < 0.05, Methods) are highlighted with * marks, with its color indicating the stronger allele. The VSMC ATAC-seq and ENCODE aortic smooth muscle cell (AoSMC) DNase tracks are shown below. (**c**) Fraction of *cis* interaction mapped to each AS profile (N=2) (**d**) ATAC-dPCR (digital PCR) on ECAD9. Significant p value was calculated by t test (N=2).

**Figure S12** Reproduction of Tri-4C using alternative digestion by CviAII. (**a**) 2D plot showing read count in 500 bp bins obtained from original Tri-4C and alternative digestion protocol. Pearson correlation coefficient r is indicated. (**b**) Loop score (log(-log(p)) comparison for all bins scored above 0 (p<0.1).

**Figure S13** (**a**) Reproducibility of Tri-HiC at various resolution for intrachromosomal contacts within 8 Mbp range (same as the distance restraint for loop calling). (**b**) Contact frequencies versus dist.nce for read pairs in indicated directions determined by Tri-HiC.

**Figure S14** Comparisons of IMR90 interaction maps between Tri-HiC (all replicates, 7.2 billion contacts), Tri-HiC (replicate #2, 1.2 billion contacts), and *in situ* HiC in at sub-kilobase resolutions in 4 chromosomal loci.

**Figure S15** (**a**) Annotation of *in situ* HiC loops. (**b**) Examples of virtual 4C derived from Tri-HiC for promoters interacting with multiple enhancers Additional annotations of Tri-HiC-identified loops. (**c**) Annotation of 1 kb Tri-HiC loops.

**Figure S16** Examples of H3K27me3-associated super-stripes (highlighted by gray bars) identified by Tri-HiC.

**Figure S17** Decomposition analysis of Tri-HiC loops. (**a**) Scatter plot showing correlations between loop fold enrichment (FE) and product of stripe fold enrichments of the two corresponding loop anchors around the loop sites. See Methods for precise definition of loop and stipe regions. (**b**) Categorial comparisons between loop and stripe fold enrichments. Loops annotated into multiple categories are only included in the leftmost category. Numbers above the bars indicate p values for the differences determined by two-sample t tests. (c) Distributions of residual loop strengths for each TF identified in pairs on both loop anchors (i.e. diagonal of (d)), in comparison with the distribution of average residual loop strength for each TF (i.e. row average). Significance of differential distribution was calculated by paired t-test. (d) Heatmap of average residual loop strengths for all 1 kb loops identified by Tri-HiC with indicated TF motif parings.

**Fig S18** Examples of loci where lncRNAs are located between the oncogene and its looped enhancer networks. For MEIS1 (green arrow), interactions with the downstream enhancers is interfered by LINC01798(red arrow) promoter, whereas for CCND1, FZD8, and VEGFA, the promoters of respective lncRNAs (LINC01488, PCAT5, and LINC01512, dotted red arrow) are not occupied with active histone markers and show no sign of interaction insulation.

**Table S1** Sequences for viewpoint-specific primers. See Online Methods for adaptor and universal primer sequences.

**Table S2** Statistics for Tri-4C and UMI-4C libraries. Total Read indicates actual sequencing depth. On Target Ratio indicates reads with matched padding sequence. Unique Read indicates yield after deduplication. Intra-TAD Ratio indicates reads falling into the same TAD as the viewpoint.

**Table S3** Sequences for gRNA and validation primers used for MLLT3 putative enhancer deletion.

**Table S4** Summary Statistics for IMR90 Tri-HiC libraries.

