## Supplementary figures and images for "Ultrafine mapping of chromosome conformation at hundred basepair resolution reveals regulatory genome architecture"

### Figure S1

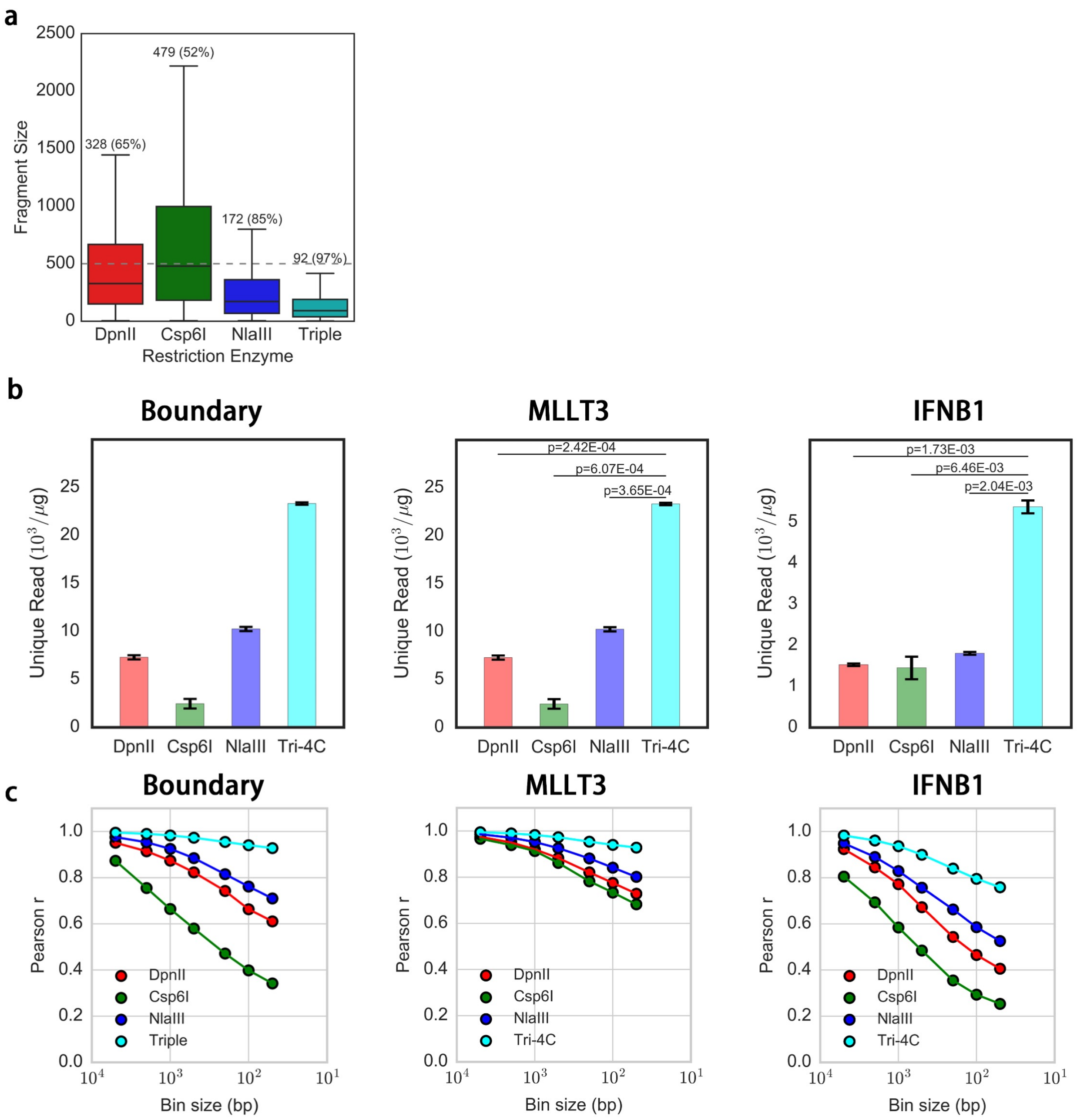

### Figure S2

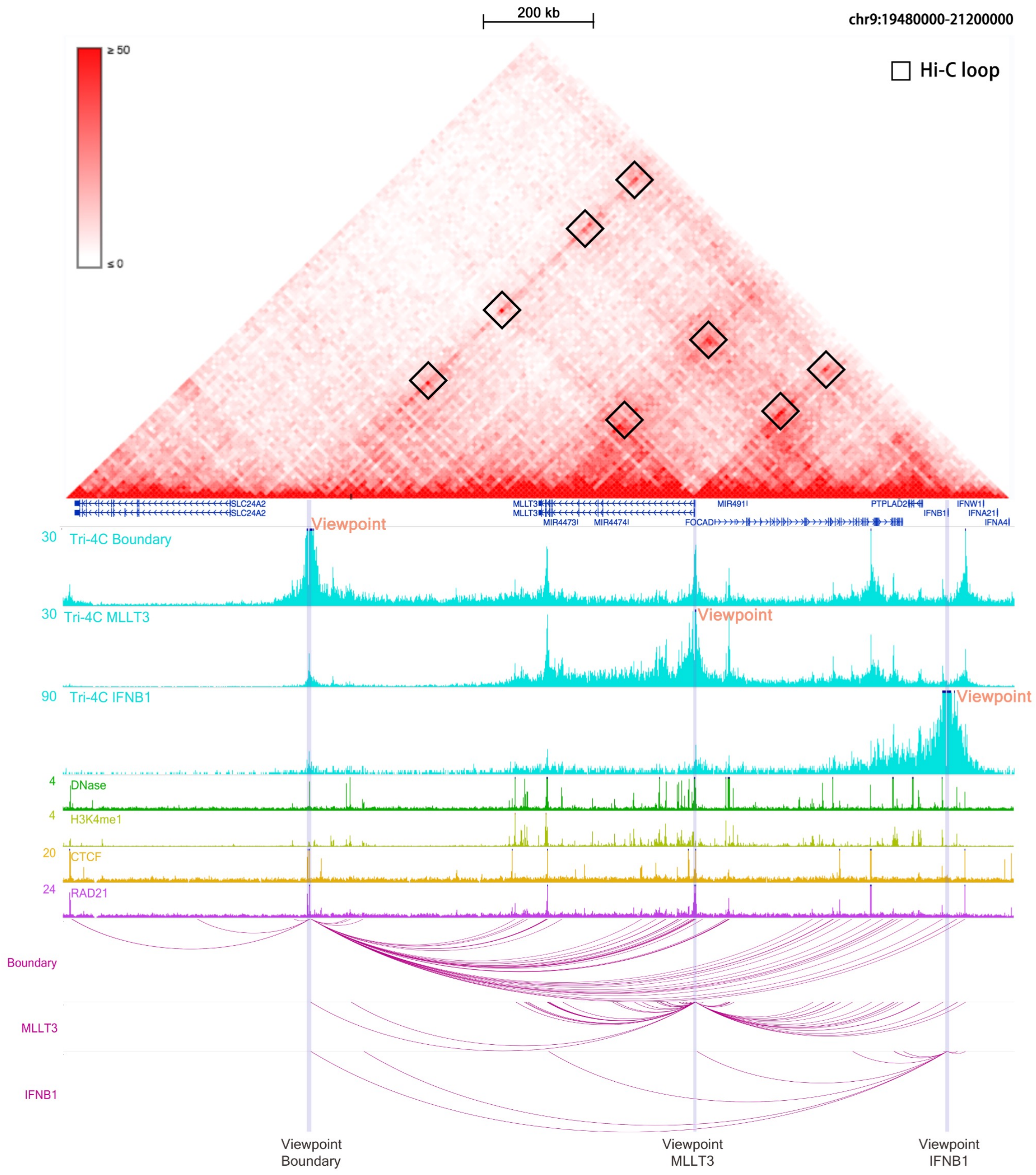

### Figure S3

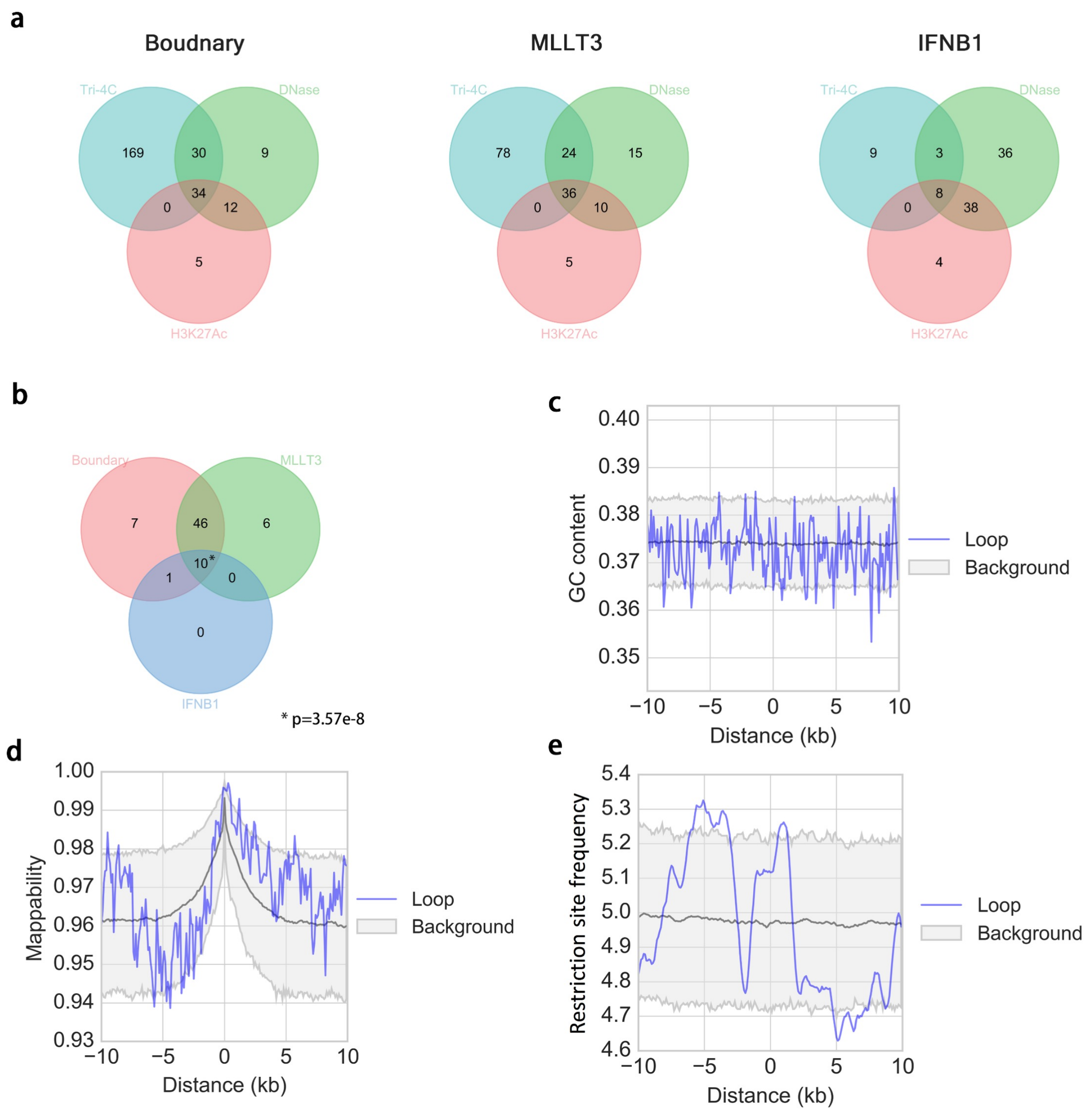

### Figure S4

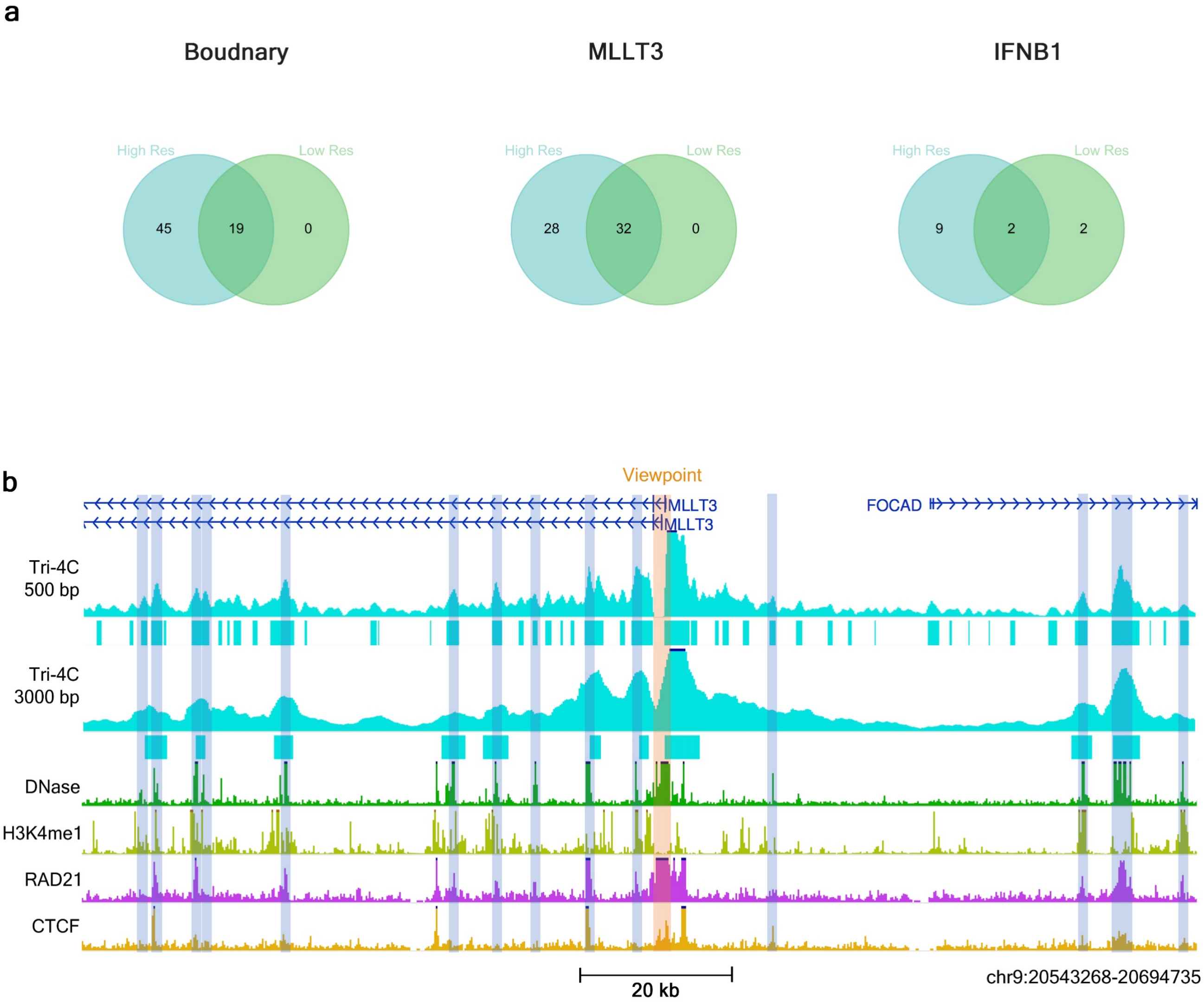

### Figure S5

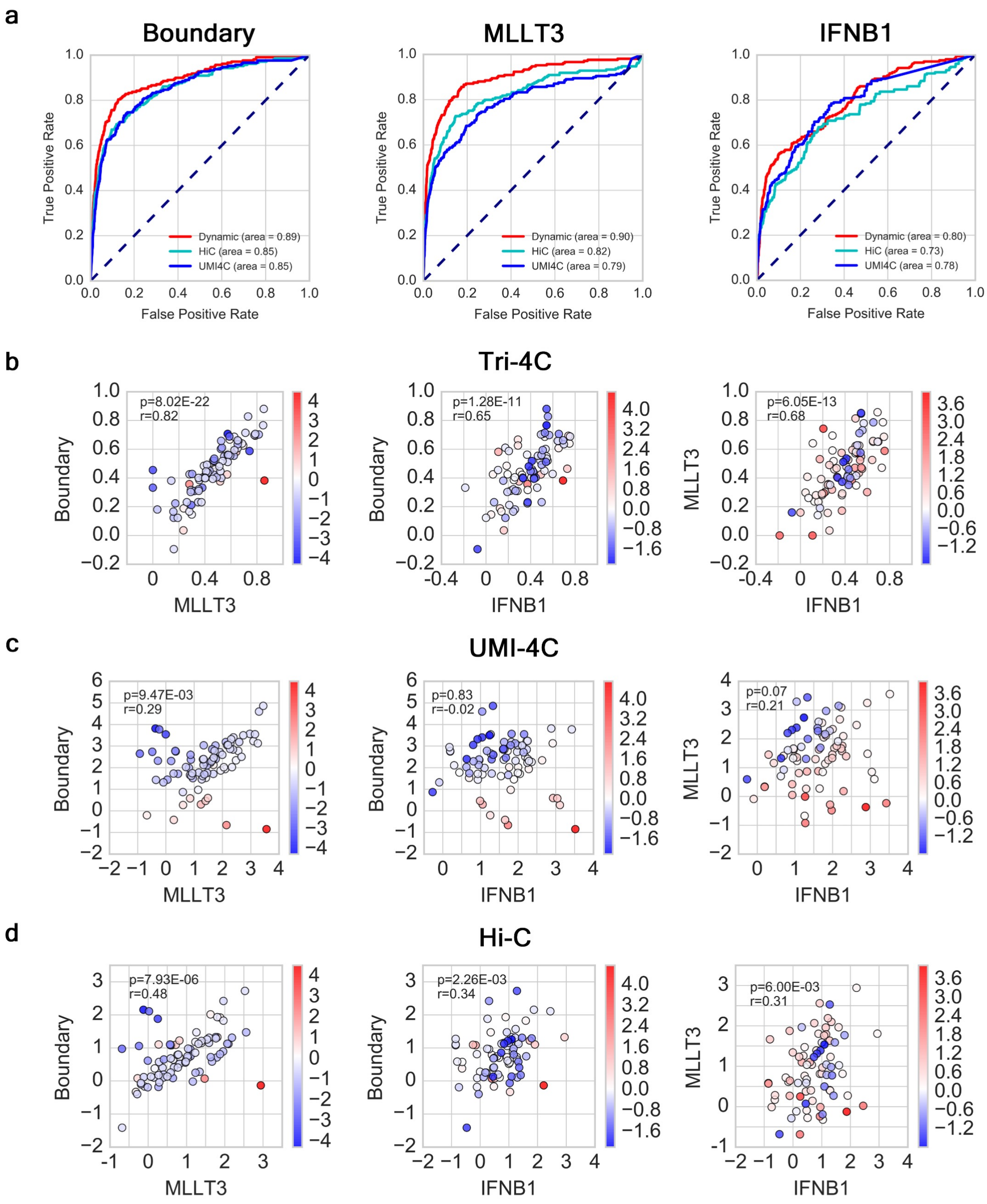

### Figure S6

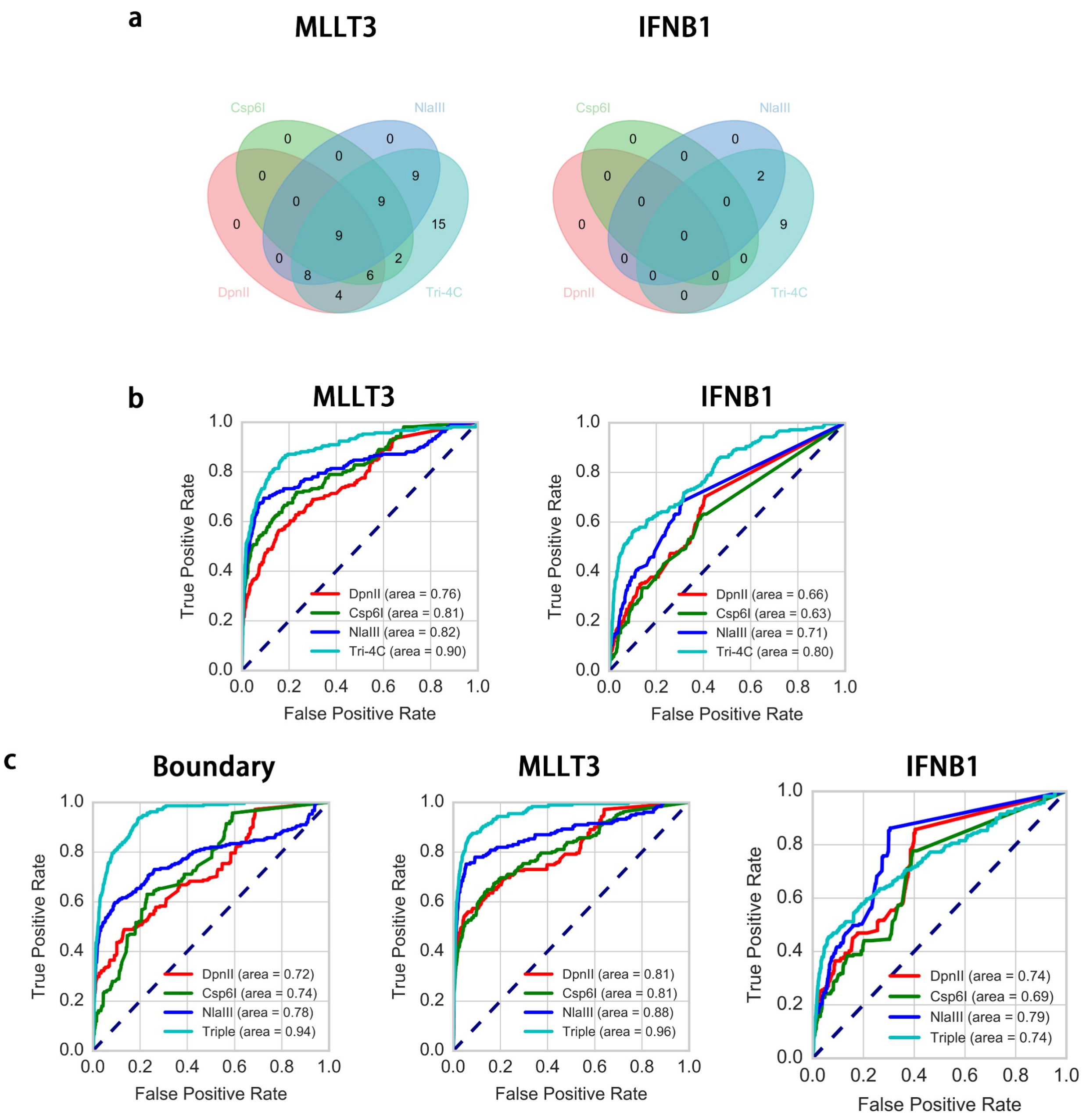

### Figure S7

**a**

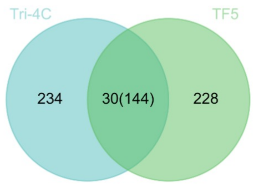


**b**

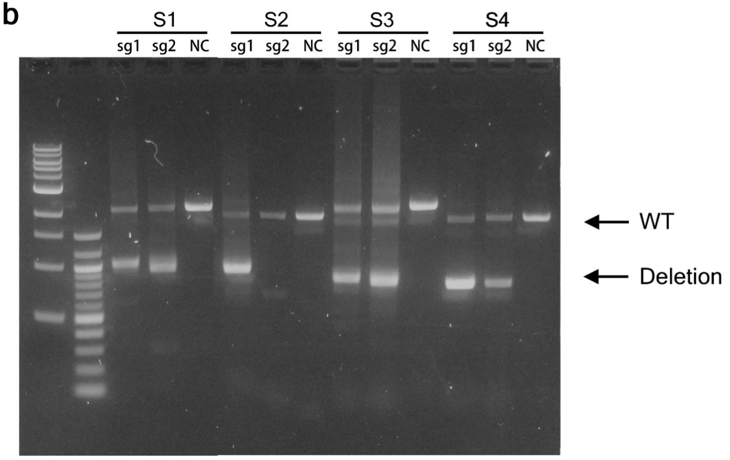

### Figure S8

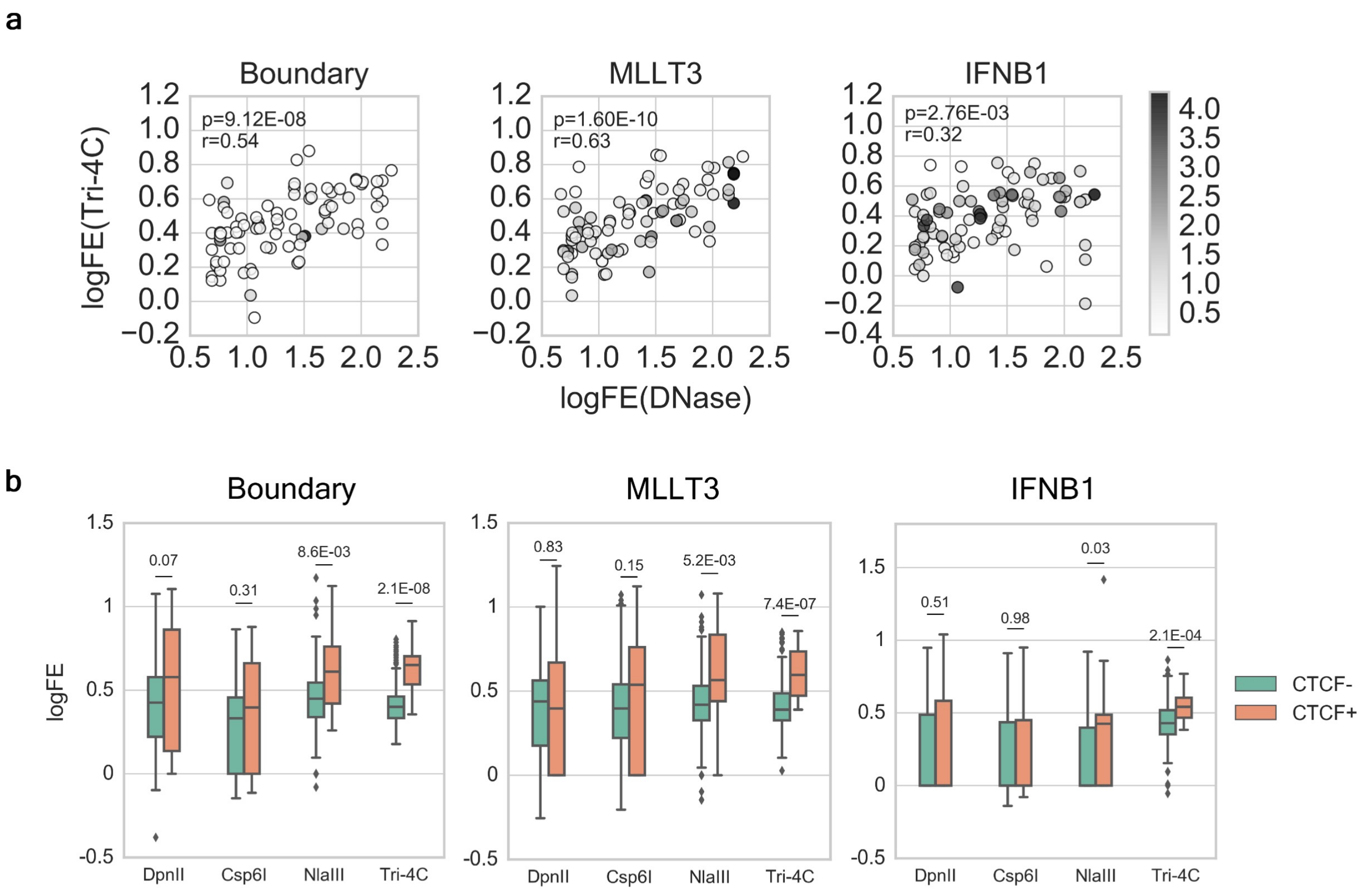

### Figure S9

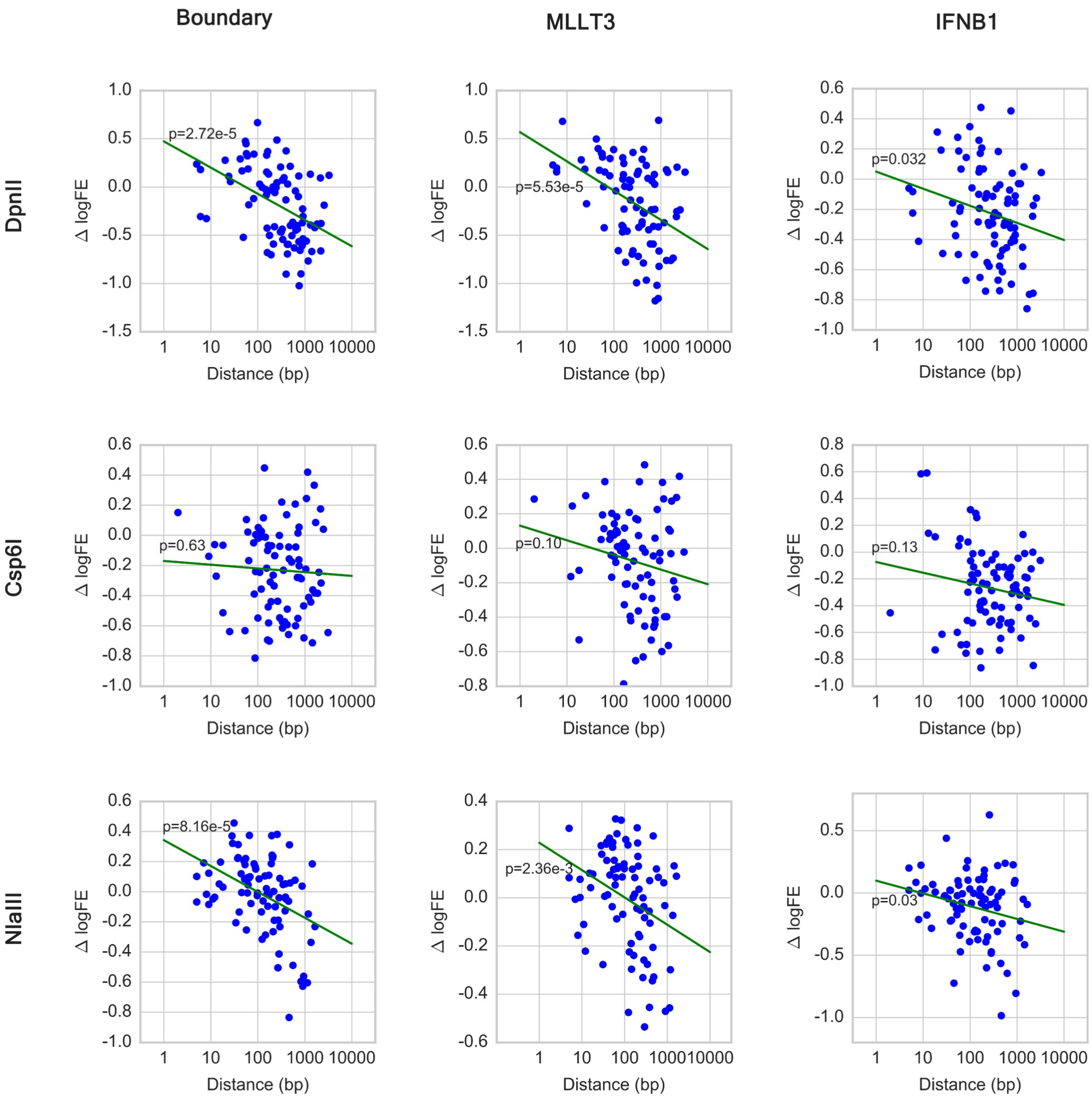

### Figure S10

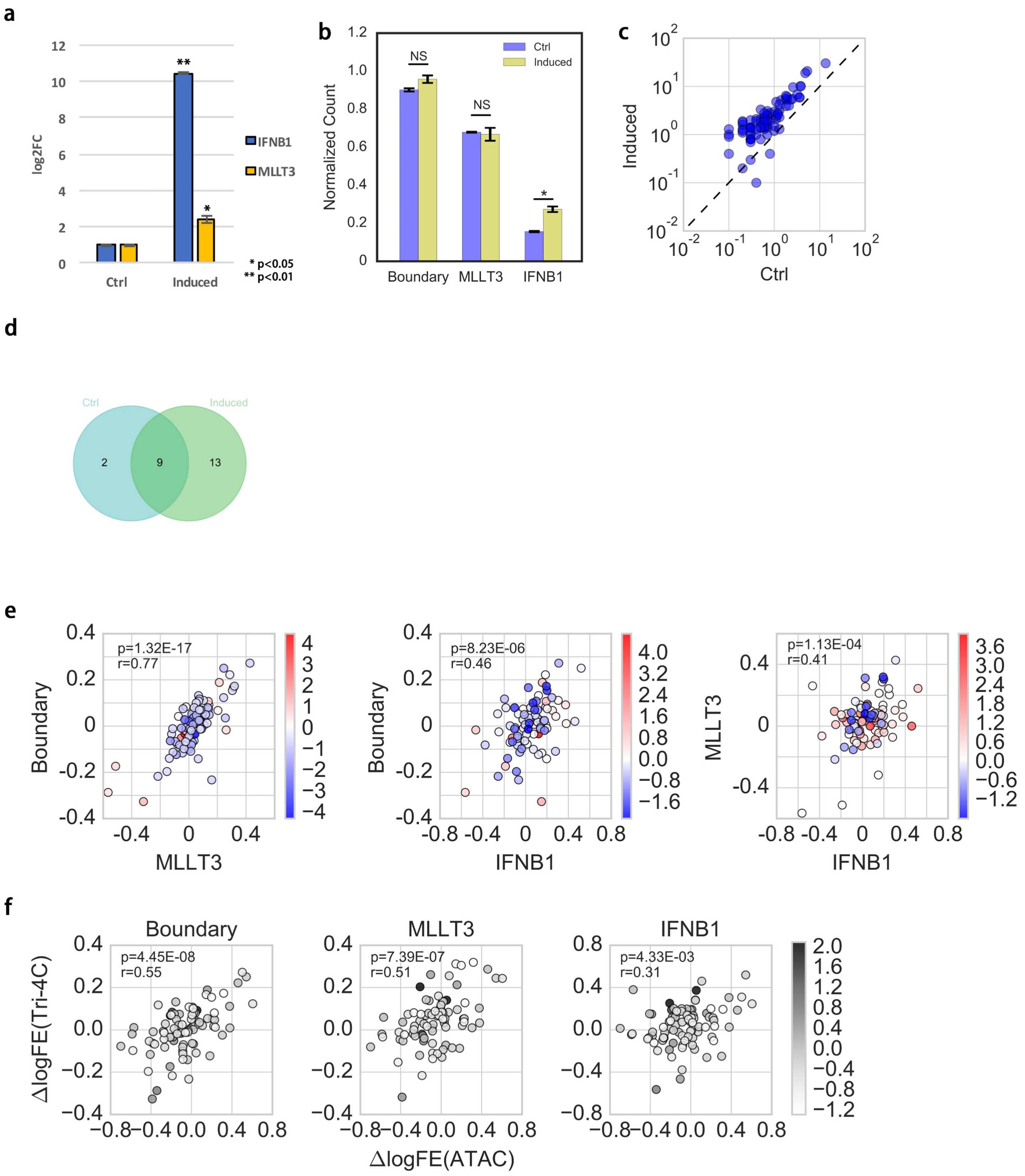

### Figure S11

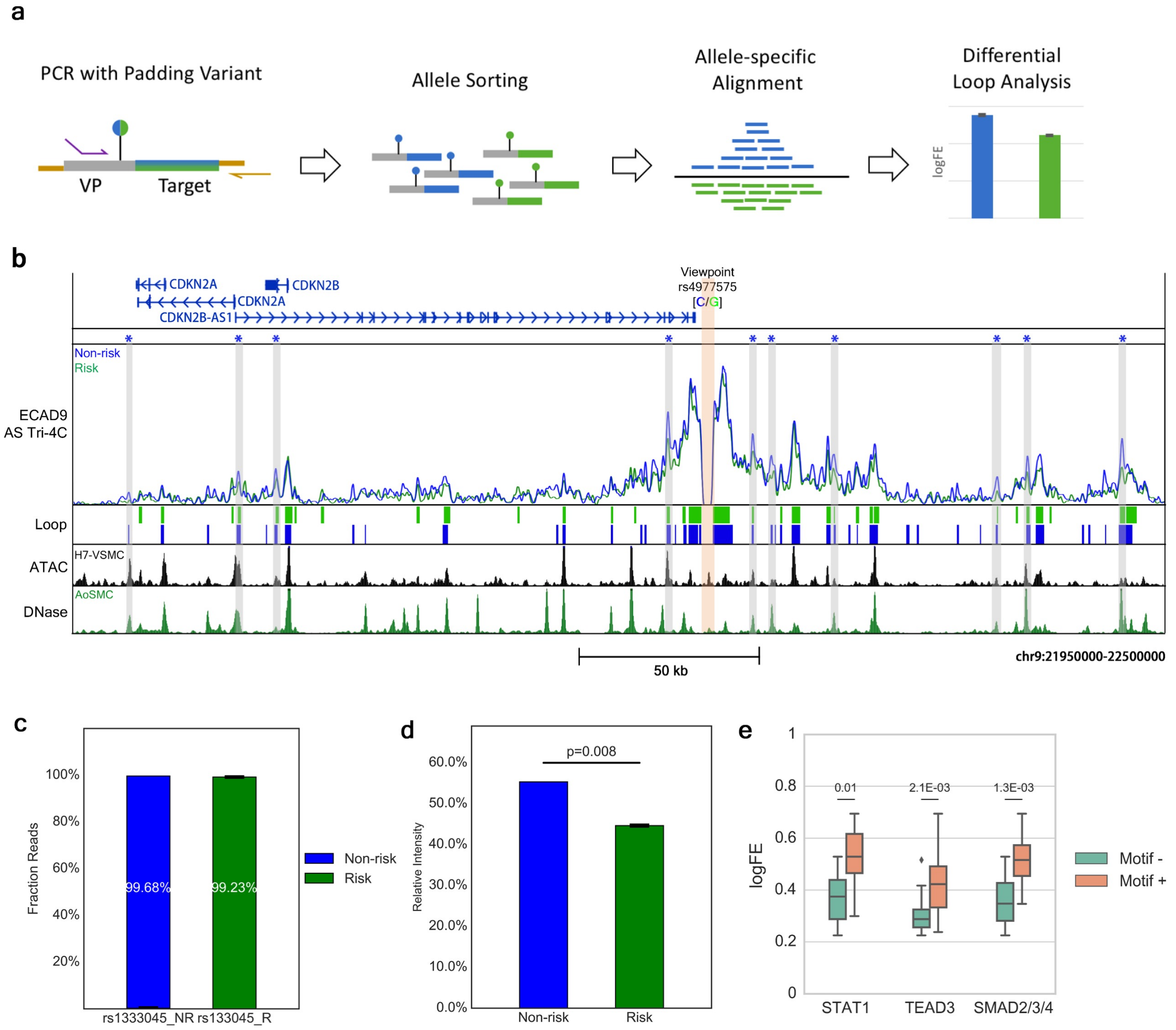

### Figure S12

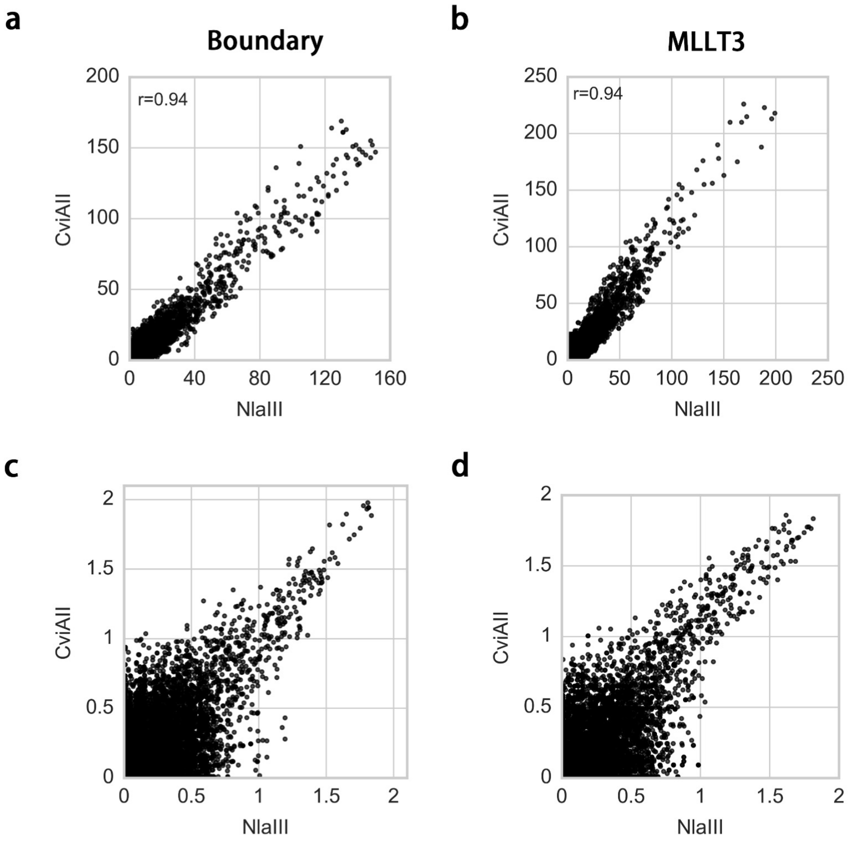

### Figure S13

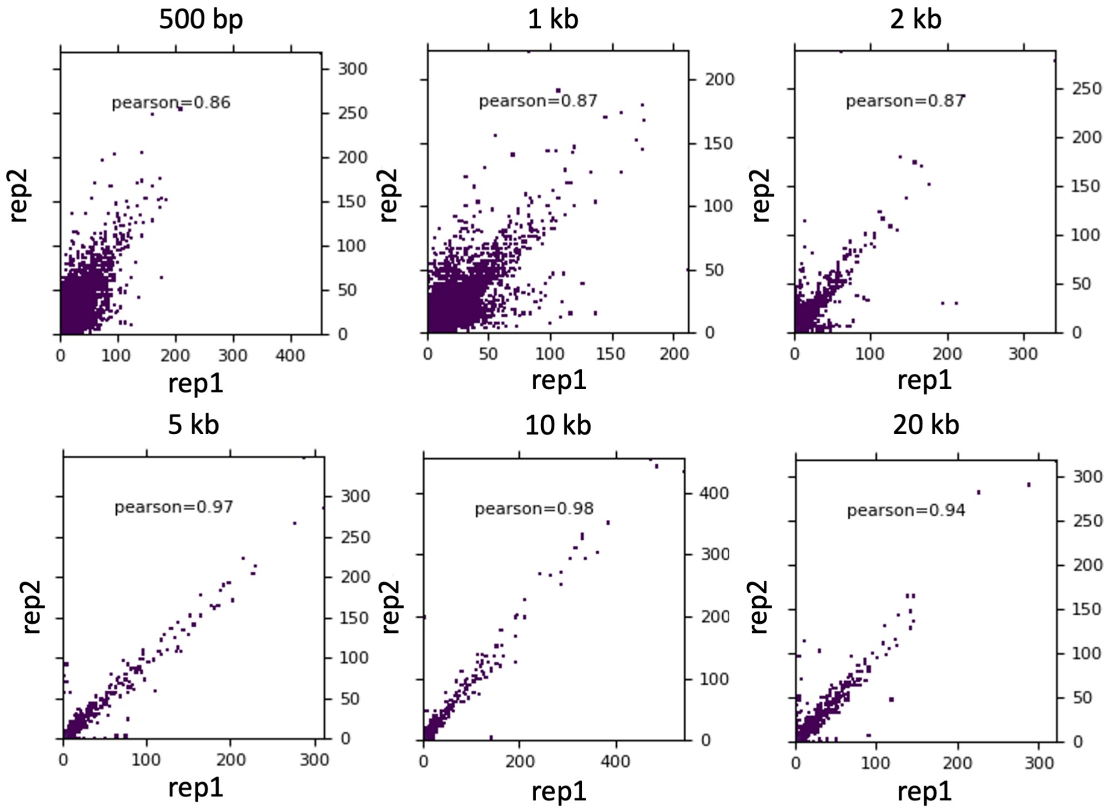

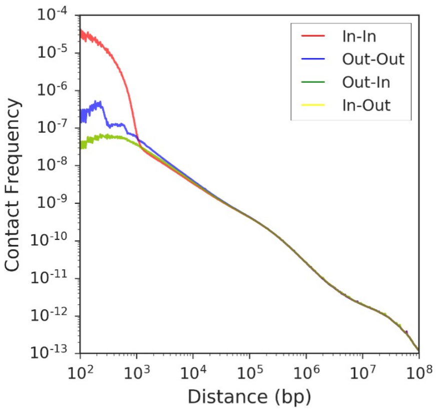

### Figure S14

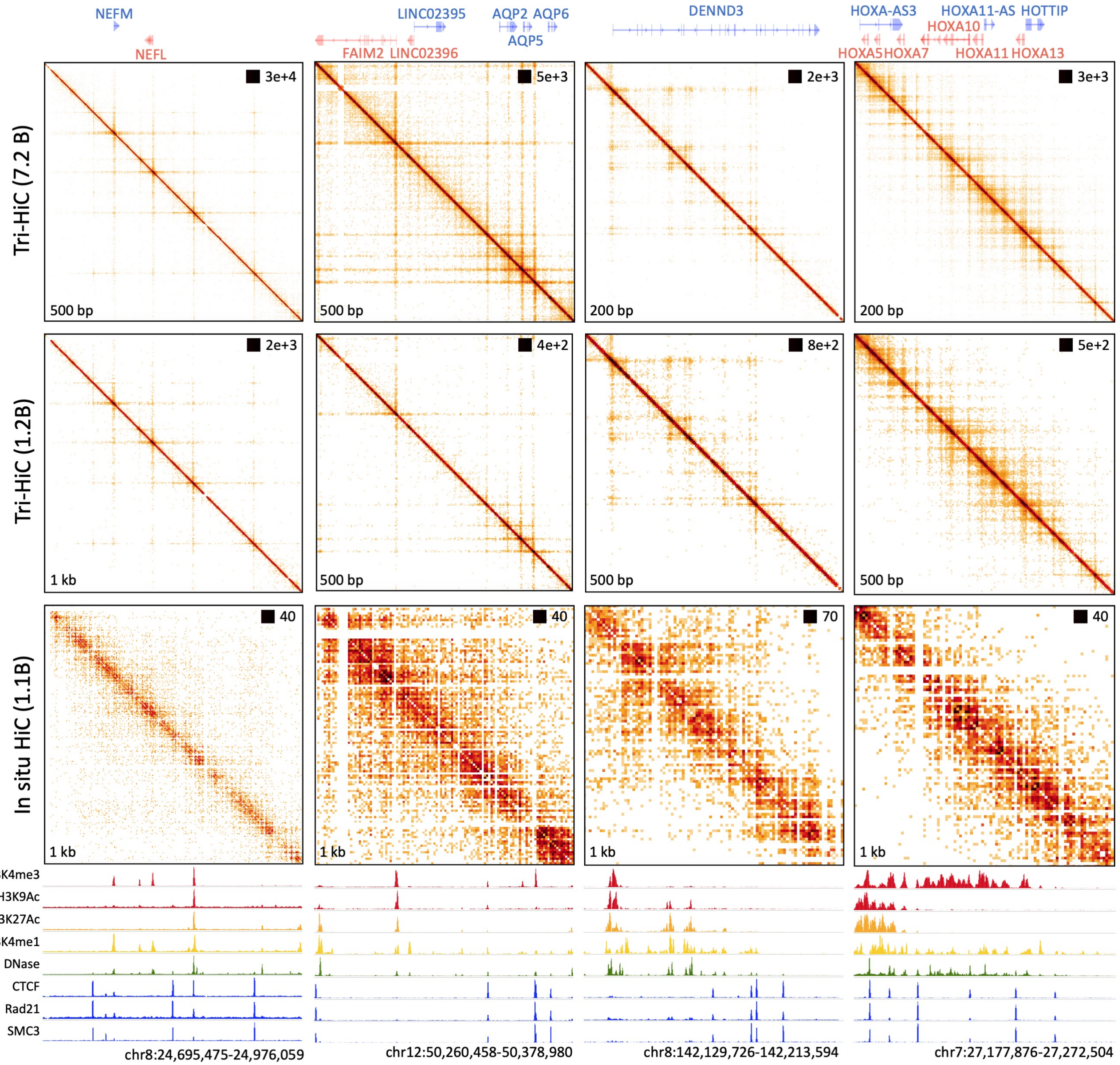

### Figure S15

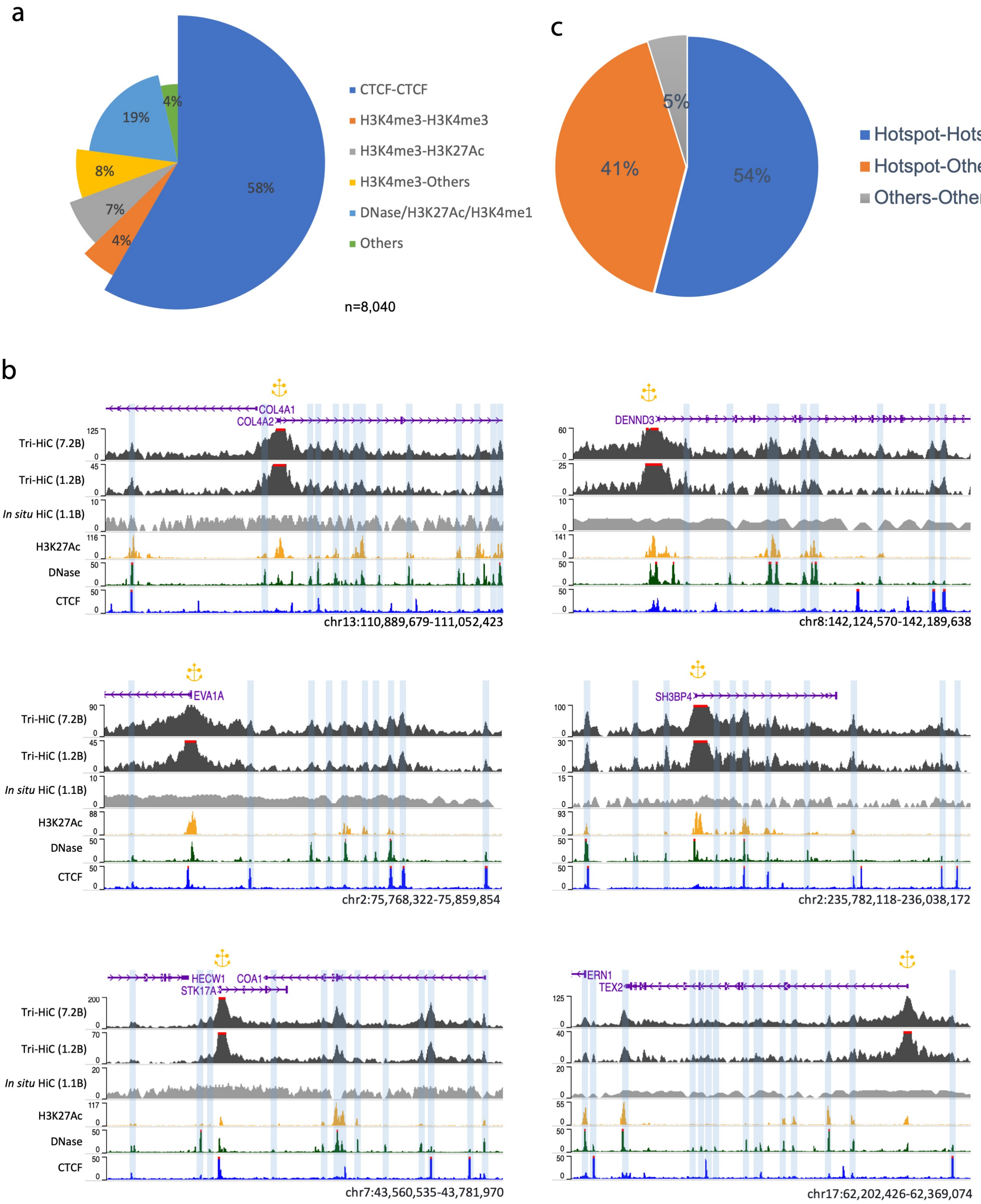

### Figure S16

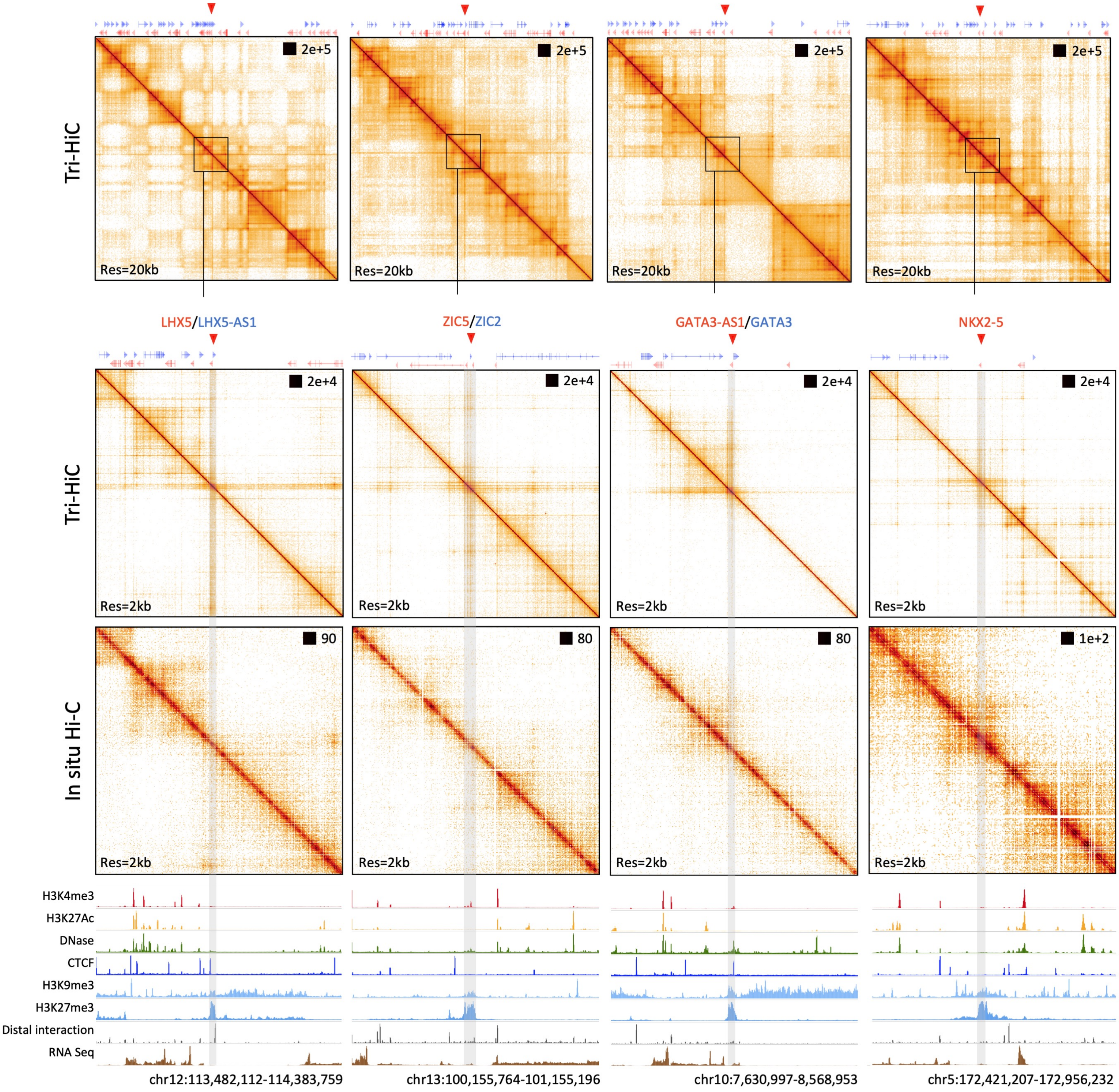

### Figure S17

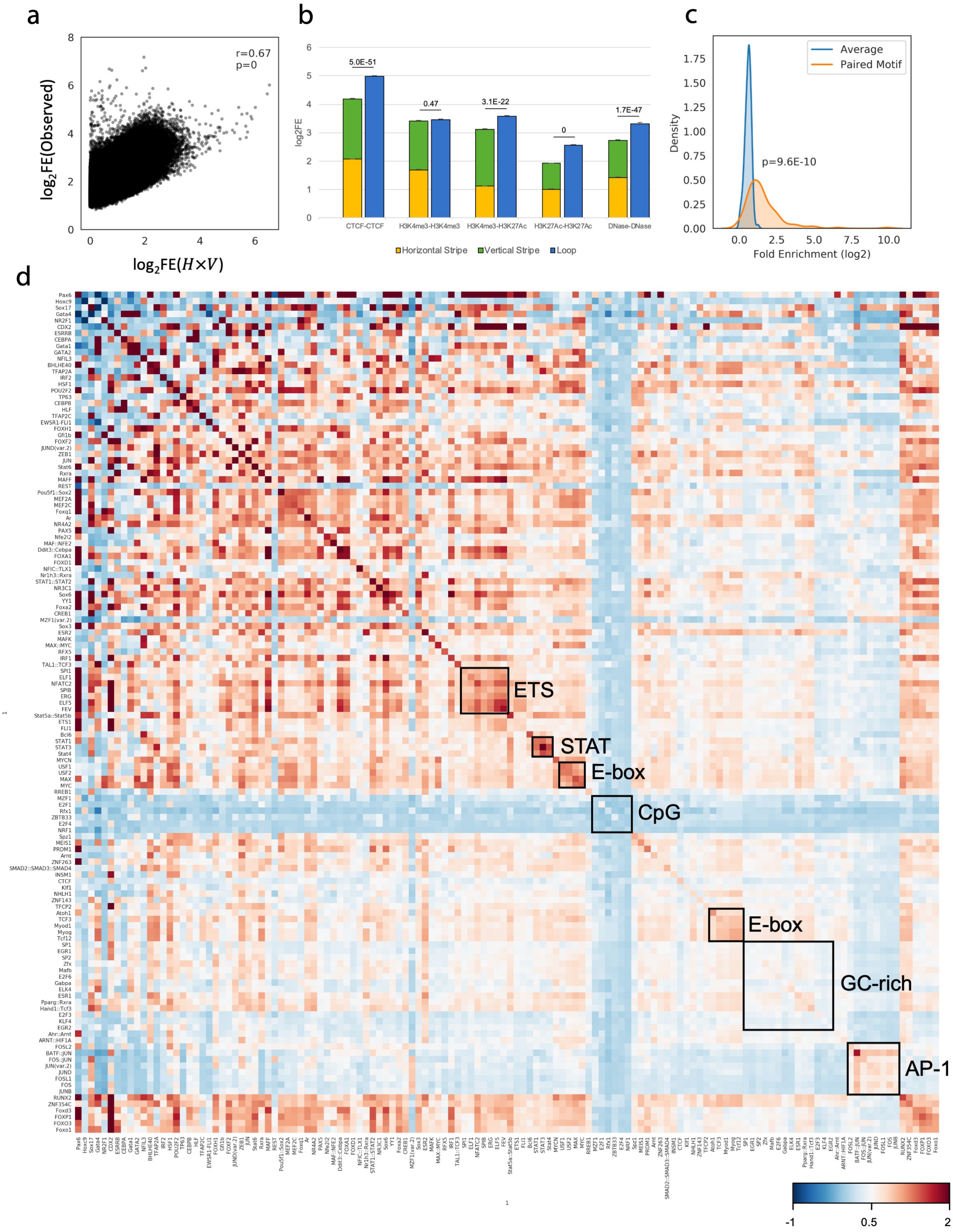

### Figure S18

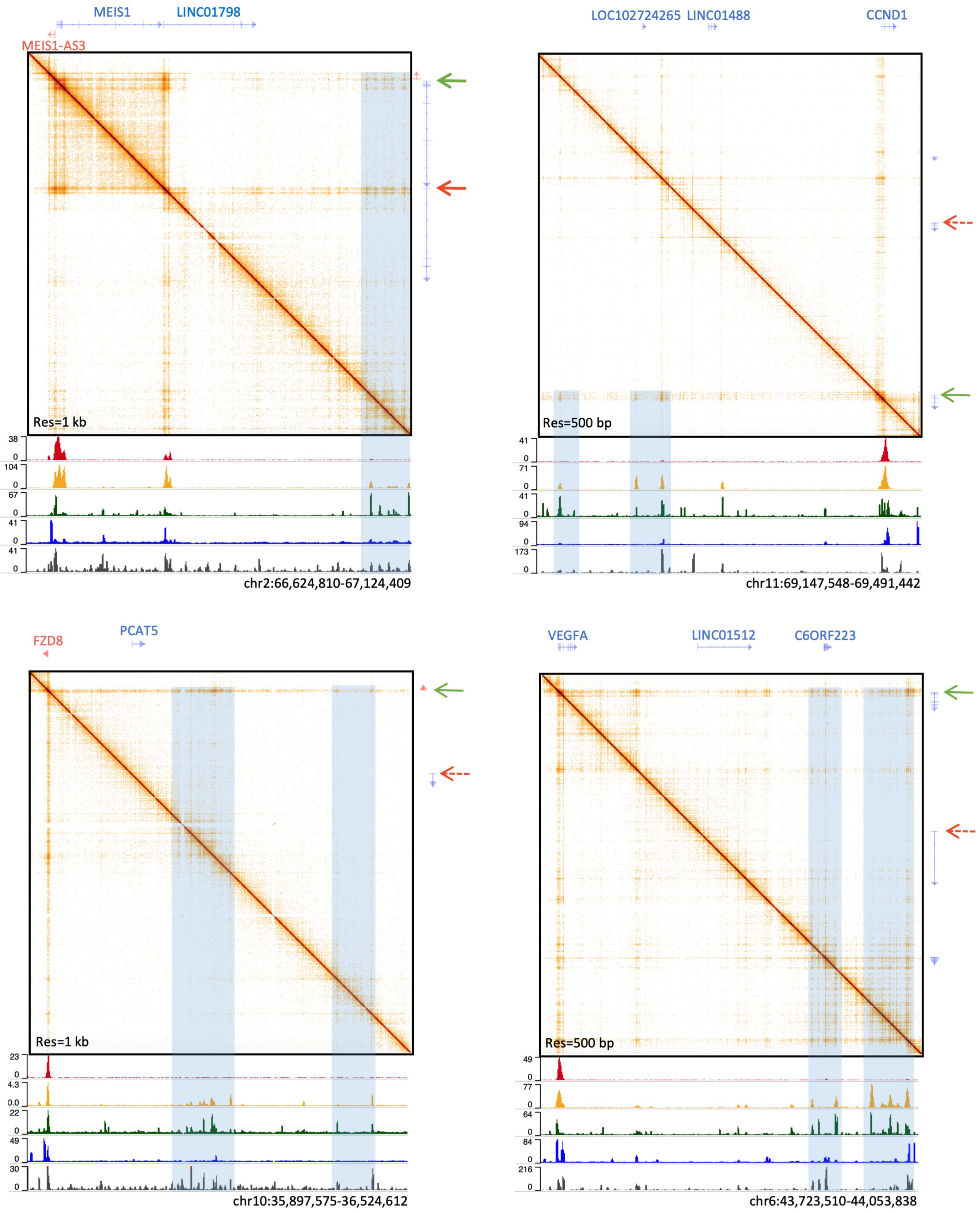
