## Supplementary material for "Ultrafine mapping of chromosome conformation at hundred basepair resolution reveals regulatory genome architecture": Table S1

| Viewpoint | Position (hg19) | Outer Primer | Inner Primer | Padding Sequence |
|---|---|---|---|---|
| Boundary-DpnII | chr9:19926377 | CACCTTTCATTCAGTACCTCCCTTCA | GAAAGCCCTGTCTCCTGTCCTTG | GACTGG |
| Boundary-Csp6I | chr9:19926333 | TTTCCTTTTCGCTCCGCTGTAGT | CCGCTGTAGTTACGTGACTCACCTTT | CATTCA |
| Boundary-NlaIII | chr9:19926164 | CGCCATAACCTCAACTCAGTCCT | CAGTCCTCTCCCGCCATCTAGT | GGT |
| Boundary-Triple | chr9:19926164 | CGCCATAACCTCAACTCAGTCCT | CAGTCCTCTCCCGCCATCTAGT | GGT |
| MLLT3-DpnII | chr9:20623064 | AACGAGCATGAAAATAGCAATGACTGA | AGCAATGACTGAGGCTGTTCTAACC | CGCAAC |
| MLLT3-Csp6I | chr9:20622883 | AGCCCTCAGCAGCCGGAGA | CGGAGAGGGGGTGTTAAATCAAGT | CCTTAG |
| MLLT3-NlaIII | chr9:20622509 | CTCCCTCCGCCCTGTGAG | CTCGGAGCCCGGGTGTCA | GGCGCCACGGCG |
| MLLT3-Triple | chr9:20622509 | CTCCCTCCGCCCTGTGAG | CTCGGAGCCCGGGTGTCA | GGCGCCACGGCG |
| IFNB1-DpnII | chr9:21077304 | GAGTGGAAATCCTAAGGAACTTTTACTTCA | TTAACAGACTTACAGGTTACCTCCGAAAC | TGAA |
| IFNB1-Csp6I | chr9:21077396 | TGAGCAGTCTGCACCTGAAAAGA | TGGGAGGATTCTGCATTACCTGA | AGGCCAAGGA |
| IFNB1-NlaIII | chr9:21077956 | GAAGTGAAAGTGGGAAATTCCTCTGA | TTCCTCTGAATAGAGAGAGGACCATC | TCATATAAATAGGCCATACC |
| IFNB1-Triple | chr9:21077956 | GAAGTGAAAGTGGGAAATTCCTCTGA | TTCCTCTGAATAGAGAGAGGACCATC | TCATATAAATAGGCCATACC |
| ECAD9[Non-risk/Risk] | chr9:22124746 | CTATCTTGAAGGCAGGCCACACT | CTTGTGTAAACAATGGTATCACATTCTAACTT | AGCTGAGAC[G/C]ACTTCTGGCCCTGA |
