## Supplementary material for "Ultrafine mapping of chromosome conformation at hundred basepair resolution reveals regulatory genome architecture": Table S2

| Viewpoint | Replication | Total Read | On Target Ratio | Unique Read | Intrachromosome Ratio | In-TAD Ratio |
|---|---|---|---|---|---|---|
| Boundary-DpnII | 1 | 9,903,873 | 0.85 | 19,803 | 0.45 | 0.65 |
| | 2 | 12,050,430 | 0.87 | 22,113 | 0.45 | 0.65 |
| Boundary-Csp6I | 1 | 3,613,316 | 0.94 | 8,632 | 0.49 | 0.65 |
| | 2 | 2,899,486 | 0.92 | 5,704 | 0.50 | 0.69 |
| Boundary-NlaIII | 1 | 2,266,904 | 0.91 | 20,003 | 0.46 | 0.61 |
| | 2 | 2,067,982 | 0.91 | 26,365 | 0.49 | 0.60 |
| Boundary-Triple_Ctrl | 1 | 6,166,334 | 0.97 | 98,426 | 0.68 | 0.68 |
| | 2 | 8,287,678 | 0.95 | 101,384 | 0.69 | 0.68 |
| Boundary-Triple_Induced | 1 | 5,648,676 | 0.93 | 102,319 | 0.65 | 0.66 |
| | 2 | 6,113,449 | 0.92 | 108,429 | 0.65 | 0.66 |
| Boundary-Triple_Alt_Dig | 1 | 3,562,529 | 0.94 | 42,037 | 0.63 | 0.73 |
| | 2 | 3,675,999 | 0.94 | 31,493 | 0.60 | 0.76 |
| MLLT3-DpnII | 1 | 4,746,406 | 0.93 | 22,643 | 0.47 | 0.73 |
| | 2 | 6,462,079 | 0.94 | 24,034 | 0.46 | 0.72 |
| MLLT3-Csp6I | 1 | 5,109,338 | 0.95 | 9,467 | 0.49 | 0.74 |
| | 2 | 4,809,153 | 0.95 | 6,271 | 0.53 | 0.78 |
| MLLT3-NlaIII | 1 | 1,613,409 | 0.89 | 32,103 | 0.52 | 0.73 |
| | 2 | 1,522,203 | 0.88 | 33,497 | 0.51 | 0.74 |
| MLLT3-Triple_Ctrl | 1 | 11,108,271 | 0.96 | 75,018 | 0.56 | 0.74 |
| | 2 | 19,623,818 | 0.96 | 74,235 | 0.57 | 0.74 |
| MLLT3-Triple_Induced | 1 | 19,488,449 | 0.93 | 68,200 | 0.55 | 0.70 |
| | 2 | 17,435,504 | 0.93 | 78,689 | 0.54 | 0.68 |
| MLLT3-Triple_Alt_Dig | 1 | 4,455,496 | 0.93 | 52,089 | 0.54 | 0.77 |
| | 2 | 3,548,450 | 0.92 | 41,754 | 0.47 | 0.80 |
| IFNB1-DpnII | 1 | 8,082,998 | 0.55 | 4,983 | 0.39 | 0.75 |
| | 2 | 9,094,900 | 0.54 | 4,791 | 0.28 | 0.72 |
| IFNB1-Csp6I | 1 | 22,362,696 | 0.29 | 3,766 | 0.23 | 0.74 |
| | 2 | 17,115,256 | 0.25 | 5,523 | 0.44 | 0.67 |
| IFNB1-NlaIII | 1 | 21,198,926 | 0.17 | 5,885 | 0.50 | 0.79 |
| | 2 | 23,438,769 | 0.17 | 5,663 | 0.42 | 0.74 |
| IFNB1-Triple_Ctrl | 1 | 12,520,527 | 0.41 | 16,682 | 0.54 | 0.72 |
| | 2 | 10,331,398 | 0.39 | 17,689 | 0.59 | 0.73 |
| IFNB1-Triple_Induced | 1 | 9,631,644 | 0.35 | 27,829 | 0.68 | 0.69 |
| | 2 | 8,253,459 | 0.35 | 32,334 | 0.67 | 0.68 |
| ECAD9 | 1 | 9,248,315 | 0.87 | 128,749 | 0.38 | 0.60 |
| | 2 | 8,074,667 | 0.82 | 89,873 | 0.41 | 0.66 |
