## Supplementary material for "Ultrafine mapping of chromosome conformation at hundred basepair resolution reveals regulatory genome architecture": Table S3

| Site | Target | Sequence | Position (hg19; chr9) |
|---|---|---|---|
| MLLT3 S1 | gRNA Left 1 | TATTTTTCAGAGTTGAAATT | 20612755 |
| | gRNA Right 1 | AACTGGGACAATCTTTTTGG | 20613816 |
| | gRNA Left 2 | ATAAAATGCTTAATCCCACC | 20612715 |
| | gRNA Right 2 | ATGTTAAATTTCTAGAAGGG | 20613878 |
| | Validation Left | TCTCAAGTGTCCTTTCCAGCTCT | |
| | Validation Right | CCTTTCCTTCCTTCCTTCTTTCA | |
| MLLT3 S2 | gRNA Left 1 | CAAGTCTATACAGCAATGAC | 20625749 |
| | gRNA Right 1 | AGGTGCTACTTATTAAATTA | 20626696 |
| | gRNA Left 2 | ACACTCTAATTACACGTTAA | 20625638 |
| | gRNA Right 2 | ATGCTTAGTGGAATAGTTCT | 20626849 |
| | Validation Left | TTGTGTACTTGACAATGTGGTTATAGAAA | |
| | Validation Right | TTGTTGAAGTTTGAGCTGTCCAA | |
| MLLT3 S3 | gRNA Left 1 | GAGGAGGATGACATAAGTGC | 20629747 |
| | gRNA Right 1 | TACCCCTCACAGACATTTTA | 20631067 |
| | gRNA Left 2 | TAAATTTTTTTATAGAGAGG | 20629851 |
| | gRNA Right 2 | TAAATATGATGACAGTGTTC | 20631240 |
| | Validation Left | CAAAAAGACGAGGATAGGTCCAG | |
| | Validation Right | GGCCTTGATTGAAGGAAAGGTTA | |
| MLLT3 S4 | gRNA Left 1 | GCTCCCCAGTTTTGCCAAGT | 20632843 |
| | gRNA Right 1 | AGATGTGGGTAATGTATGGT | 20633966 |
| | gRNA Left 2 | CGGGTATAAGCAAGCCCACT | 20632845 |
| | gRNA Right 2 | GAACAGATGTGGGTAATGTA | 20633962 |
| | Validation Left | GGGCATTTGTCATTACACAGGAT | |
| | Validation Right | TCCTGTACCTGTCTCAATGATGC | |
| GFP | Negative Ctrl | GAAGTTCGAGGGCGACACCC | |
