## Supplementary material for "Ultrafine mapping of chromosome conformation at hundred basepair resolution reveals regulatory genome architecture": Table S4

| | Rep 1 | Rep 2 | Rep 3 | Rep 4 | Rep 5 |
|---|---|---|---|---|---|
| Raw reads | 1,043,541,346 | 2,074,938,244 | 3,196,905,866 | 3,071,890,673 | 2,965,478,312 |
| Mapped Contacts | 889,775,906 | 1,765,192,278 | 2,717,276,890 | 2,606,887,017 | 2,515,356,700 |
| Unique Contacts | 636,603,108 | 1,242,385,471 | 1,808,577,192 | 1,789,041,704 | 1,701,377,395 |
| Cis Contacts | 592,135,524 | 1,153,435,774 | 1,676,938,486 | 1,657,679,775 | 1,578,679,264 |
| Trans Contacts | 44,467,584 | 88,949,697 | 131,638,706 | 131,361,929 | 122,698,131 |
